# *De novo* designed bright, hyperstable rhodamine binders for fluorescence microscopy

**DOI:** 10.1101/2025.06.24.661379

**Authors:** Yuda Chen, Klaus Yserentant, Kibeom Hong, Yiming Kuang, Arghya Bhowmick, Arthur Charles-Orszag, Samuel J. Lord, Lei Lu, Kaipeng Hou, Samuel I. Mann, Jonathan B. Grimm, Luke D. Lavis, R. Dyche Mullins, William F. DeGrado, Bo Huang

## Abstract

*De novo* protein design has emerged as a powerful strategy with the promise to create new tools. The practical performance of designed fluorophore binders, however, has remained far from meeting fluorescence microscopy demands. Here, we design *de novo* Rhodamine Binder (Rhobin) tags that combine ideal properties including size, brightness, and now adding hyperstability. Rhobin allows live and fixed cell imaging of a wide range of subcellular targets in mammalian cells. Its reversible fluorophore binding further enables live super-resolution STED microscopy with low photobleaching, as well as PAINT-type single-molecule localization microscopy. We showcase Rhobin in the extremophile *Sulfolobus acidocaldarius* living at 75°C, an application previously inaccessible by existing tags. Rhobin will serve as the basis for a new class of live cell fluorescent tags and biosensors.

## Introduction

Protein engineering, in which one uses natural proteins as starting points for protein design, has provided a wealth of new tools that have transformed our understanding of biology and therapeutics. For example, fluorescence microscopy has been revolutionized by protein tags that create or bind a fluorescent species. Fluorescent proteins (FPs) were engineered extensively from natural proteins that have the capability to spontaneously oxidize to form chromophores (*1*). Chromophore-binding proteins have also been engineered into tags, overcoming the slow maturation and limited spectral range of FPs. These types of tags typically exhibit low brightness and/or photostability, however, due to the intrinsic limitations of their fluorophore (*2*, *3*). An alternative strategy is to engineer enzymes into self-labeling tags such as HaloTag (*4*) and SNAP-tag (*5*), which recognize distinct ligand moieties that can be attached to a variety of fluorophores. Despite these advances, engineering natural proteins can be time-consuming and suffer limitations. For example, extant, bright tags are incompatible with organisms that live under extreme conditions, such as thermophiles, due to the instability of natural protein scaffolds at elevated temperatures. By contrast, *de novo* design circumvents evolutionary imprints of a starting natural protein, allowing one to tailor new tools with the desired features in the initial design (*6–12*).

We set out to design fluorescent tags that bind small-molecule fluorophores with the following characteristics: 1) high brightness and photostability with single-molecule level sensitivity 2) spectral diversity, particularly in the far-red range, 3) fast labeling kinetics and high labeling efficiency, 4) smaller size than FPs and HaloTag, 5) high thermal stability, and 6) orthogonality to existing tags for multiplexed measurements. Although *de novo* proteins binding dyes such as zinc-porphyrin and FP chromophore analogs have been reported (*13–18*), they have not been adopted by the microscopy community as fluorescent tags due to the use of suboptimal fluorophores. We focused on the development of rhodamine-binding proteins, which are perhaps the most widely used class of small-molecule fluorophore and have been optimized for superior brightness and photostability. Rhodamines exist in equilibrium between a nonfluorescent lactone and a fluorescent zwitterion, and the equilibrium constant, *K*_L–Z_, dictates important properties like bioavailability and fluorogenicity (*19–21*). Here, we use *de novo* protein design to achieve all of our design goals, yielding highly thermostable proteins that bind different rhodamine dyes and creating a new class of fluorescent tags useful for cellular microscopy.

### *De Novo* Design of Rhobin

Our lab and others have designed many ligand-binding proteins based on the four-helical bundle scaffold (*22–27*), which exhibits high designability and stability. The binding cavities of previously parametrically generated four-helical bundles are not suitable to fit the non-planar rhodamine structures, however, which prevents stabilizing the dye in the fluorescent, zwitterionic form, which has a carboxyphenyl group orthogonal to the plane of the fluorophore. To create a new binding scaffold while still leveraging the stability of helical bundles, we used knowledge-guided RFdiffusion (*28*) to generate the protein scaffold (Fig. 1A, fig. S1): A shared folding core of previously designed four-helical bundle proteins was used as a starting point to guide the backbone generation for improved success rate and stability (*29*). The binding site design was anchored by fixing a statistically favorable location of an Arg residue intended to target the carboxylate group of the rhodamine ligand in the zwitterionic form (*24*). To form a binding pocket with sufficient space for rhodamines with diverse substituents, we started with a hypothetical rhodamine with bulkier modifications than those practically used. We selected one scaffold with high compactness and ligand buried surface that breaks a helix near the middle of the bundle and places a fifth helix at an acute angle relative to the long axis of the bundle to create the desired cavity (fig. S2). This scaffold has a molecular weight of ∼16 kDa, less than half the size of HaloTag7.

**Fig. 1.**
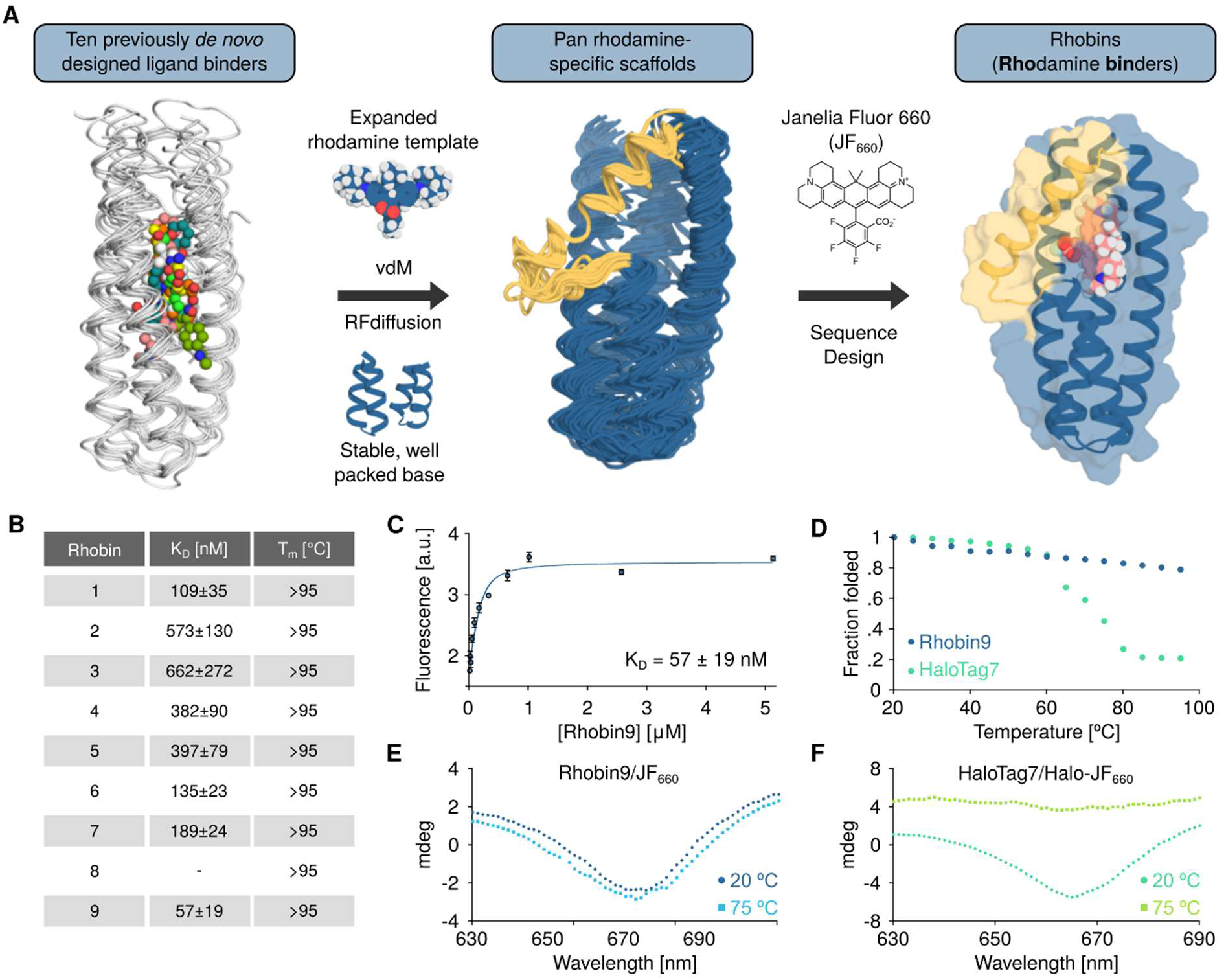
*De Novo* design of rhodamine binders. (**A**) Overall design strategy. Rhodamine-specific protein scaffolds were generated using the stably packed core of a previously *de novo* designed ligand binder, an expanded rhodamine template, and statistically preferred polar interactions at the active site as inputs to guide RFdiffusion. The selected scaffolds were then used for sequence design toward JF_660_ binding using LigandMPNN and Rosetta FastRelax. (**B**) Binding constants (*K_D_*) of all nine designs (Rhobin1 to Rhobin9) to JF_660_, measured by fluorescence titration, along with their melting temperatures determined by circular dichroism. *K_D_* is shown as mean and standard error of the mean from three independent measurements (**C**) A titration of Rhobin9 into JF_660_ shows a single-site binding isotherm. Error bars represent standard deviations (SD) from three independent measurements. (**D**) The fraction of folded protein was monitored by CD at 222 nm for both Rhobin9 and HaloTag7. (**E**) The induced CD signal of JF_660_ at 660 nm was persistent at 75 °C for the Rhobin9/JF_660_ interaction. (**F**) The induced CD signal of Halo-JF_660_ bound to HaloTag7 visible at 20 °C was diminished at 75 °C.

For sequence design, we selected the recently developed carborhodamine Janelia Fluor 660 (JF_660_) with far-red emission and excellent extinction coefficient, quantum yield, and photostability as the target (*30*). The selected scaffold was designed with an iterative process of LigandMPNN (*31*) and Rosetta FastRelax (*32*) (See SI for details). Nine designed proteins (Rhobin1 to Rhobin9) were picked after structure prediction with RaptorX-single (*33*) and sequence filtration. All nine designs agree well with structure predictions within sub-Å RMSD, including ligand-free structures from AF3 prediction and ligand-bound structures from Chai1 prediction (*34*) (fig. S3 and table S1). All nine Rhobins expressed well in *E. coli*, and were monomeric as assessed by size exclusion chromatography (fig. S4). For eight out of the nine designs, the addition of purified Rhobin to JF_660_ solution led to fluorescence enhancement, indicative of forming a non-covalent complex (fig. S5). Measuring this fluorogenic response showed that these eight Rhobins all bind JF_660_ with nanomolar affinity, with the most potent Rhobin9 at 57 nM (Figs. 1B,C). The helical bundle folding core gives Rhobin outstanding thermostability (figs. S6,7). All nine designs have melting temperatures (T_m_) of over 95 °C, compared with ∼75 °C for HaloTag7 (Fig. 1D). The non-covalent Rhobin9/JF_660_ interaction is stable at 75 °C as shown by induced circular dichroism (CD) measurements, whereas the induced CD signal of covalently attached JF_660_ to HaloTag7 disappeared at this temperature (Fig. 1E).

### Benchmarking of Rhobin in Mammalian Cells

To evaluate the performance of Rhobins in mammalian cells, we created calibration constructs where the sequence of histone H2B was fused to the sequence of both the candidate tag and mNeonGreen (as an expression level marker), with a nuclear localized mTagBFP separated by an internal ribosomal entry site (IRES) as the segmentation marker (Fig. 2A). Transient expression of the calibration construct encoding Rhobin9 showed specific nuclear signal in live human U2OS cells within minutes after addition of 100 nM JF_660_ to the culture media, contrasting the non-binding negative control (Fig. 2B). Across the range of expression levels for individual cells, signals from JF_660_ and mNeonGreen channels show strong linear relationship (R^2^ 0.83-0.87, Fig. 2C), allowing us to use the slope of the linear fit to quantify the normalized brightnesses of different binder/dye combinations. Extending this assay to other Rhobins showed trends largely consistent with *in vitro* measurements, with Rhobin9 generating the strongest on-target signal, 229× above the dye background, followed by Rhobin6 (Fig. 2D, fig. S8). We also observed efficient expression of all Rhobins except Rhobin5 (Fig. 2D, fig. S9).

**Fig. 2.**
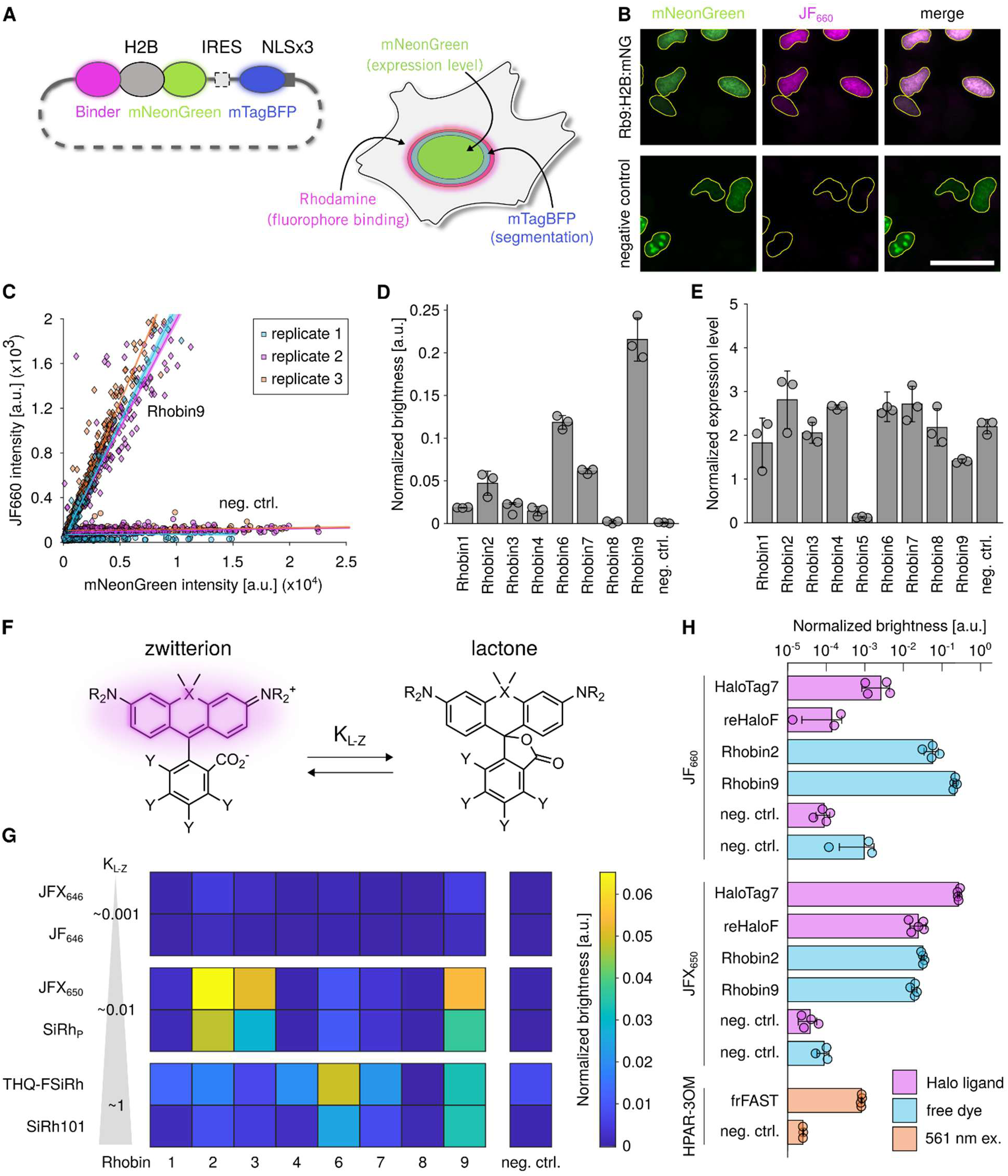
Benchmarking of Rhobin variants in mammalian cells. (**A**) Screening strategy to evaluate fluorophore binders in mammalian cells. (**B**) Widefield fluorescence images show specific Rhobin9 labeling with 100 nM JF_660_ without washing compared to the negative control of the similarly sized frFAST, which does not bind JF_660_. Yellow lines: nuclear segmentation based on mTagBFP-NLS signal. Scale bar: 50 μm. (**C**) Signal from JF_660_ binding to Rhobin9 scales linearly with expression level, showing per-cell mean nuclear JF_660_ and mNeonGreen intensities and linear fits from three independent replicates. (**D, E**) Rhobin variants efficiently bind JF_660_ (D) and express well (E) in U2OS cells. The normalized brightnesses and normalized expression levels were determined by the slope of linear fits to the per-cell JF_660_ to mNeonGreen (D) and mNeonGreen to mTagBFP (E) mean nuclear intensities. See figs. S8, S9 for individual scatter plots. frFAST as the negative control. Showing individual data points and mean ± SD. (**F**) Rhodamines exist in an equilibrium (*K*_L-Z_) between a fluorescent zwitterionic (left) and a non-fluorescent lactone (right) form. (**G**) Rhobin variants exhibit distinct brightness profiles across rhodamine derivatives. See table S2 for numerical values and fig. S10 for dye structures. (**H**) The ligand-dependent brightnesses of Rhobin, HaloTag7, and reHaloF. Calibration constructs were transiently expressed in U2OS cells, labeled with their respective ligands: JFX_650_/JF_660_ for Rhobin: Halo-JFX_650_/Halo-JF_660_, for HaloTag7 and reHaloF, HPAR-3OM for frFAST, and imaged under no-wash conditions. The normalized brightness for each condition was obtained from linear fits as described above. Showing individual data points and mean ± SD. See table. S2 for numerical values and table S3 for imaging conditions.

Although JF_660_ produces strong fluorescence upon Rhobin binding, it primarily exists in the fluorescent zwitterionic form (*K*_L-Z_ 7.55) (*30*), which could lead to appreciable background from unbound dye molecules. Other fluorophores, especially Si-rhodamines, exhibit lower *K*_L–Z_ values that can yield fluorogenic ligands that exhibit lower background. We examined the binding of three pairs of far-red emitting Si-rhodamines that are structurally similar to JF_660_ but have a range of *K*_L-Z_ values (*30*, *35*, *36*) (Fig. 2F, fig. S10). Two dyes with high *K*_L-Z_ (∼1, THQ-FSiRh, SiRh101) showed normalized brightness profiles similar to that of JF_660_, which is expected due to the propensity of these dyes to adopt the fluorescent form and the analogous structures containing fused julolidine or tetrahydroquinoline ring systems. Two dyes containing pyrrolidine auxochromes, SiRh_P_ and its deuterated version JFX_650_ have intermediate *K*_L-Z_ values (∼0.01) and show a different profile, binding to Rhobin2 and Rhobin3 to produce the strongest signal, followed by binding to Rhobin9. Finally, we tested the azetidine-containing JF_646_ and its deuterated version JFX_646_. They have low *K*_L-Z_ values (∼0.001) and show weak but specific signals, with the normalized brightness profile across Rhobin variants closer to those of SiRh_P_/JFX_650_. These results show differences across the different Rhobin variants. Rhobin9 is a relatively promiscuous binder, whereas Rhobin2, Rhobin3, and Rhobin6 show increased selectivity for different rhodamine derivatives. Based on these results, we select Rhobin2 and JFX_650_/SiRh_P_ as our fluorogenic labeling system in addition to the Rhobin9/JF_660_ system.

We then compared Rhobin2 and Rhobin9 against established fluorophore binding protein tags, including HaloTag7 (*4*), the non-covalent ligand binder reHaloF (*37*), and the dye-binding frFAST (*38*) that transiently binds the HPAR-3OM fluorophore. The normalized brightness of Rhobin2/9 with JF_660_ is substantially higher than HaloTag7/Halo-JF_660_ or reHaloF/Halo-JF_660_ (Fig. 2G). Rhobin2/9 with JFX_650_ is similar to reHaloF/Halo-JFX_650_ and lower than the HaloTag7/Halo-JFX_650_ conjugate. The Rhobin system was substantially brighter than frFAST, which exhibited <1% of the signal generated by the Rhobin9/JF_660_ system and <3% of the Rhobin2/JFX_650_ pair with blue-shifted excitation and lower photostability.

### Cellular Microscopy Using Rhobin

To assess the versatility of Rhobin for protein tagging in mammalian cells, we established U2OS cell lines stably expressing various Rhobin2 fusion proteins by integrating the constructs into the *Pa01* attB landing pad (*39*) at the *AAVS1* safe harbor locus (Fig. 3A). These cells were isolated by fluorescence-activated cell sorting (FACS) after labeling with SiRh_p_. The cells maintained normal proliferation across multiple passages, indicating that sustained expression of Rhobin2 fusions does not disrupt cellular proliferation. Rhobin2/SiRh_p_ robustly labeled a wide range of subcellular compartments—including the nucleus (histone H2B), nucleoli (NPM1 for granular component; FBL for dense fibrillar component), nuclear lamina (LMNA), actin filaments (LifeAct), endosomes (Rab5a), clathrin-coated vesicles (CLTB), endoplasmic reticulum (ER, SEC61B), Golgi apparatus (Giantin), and the mitochondrial outer membrane (TOMM20), with expected localization and morphology in both live and fixed cells (Fig. 3B). These results show that Rhobin does not perturb native protein localization and is broadly compatible with diverse subcellular structures as a fluorescent imaging tag.

**Fig. 3.**
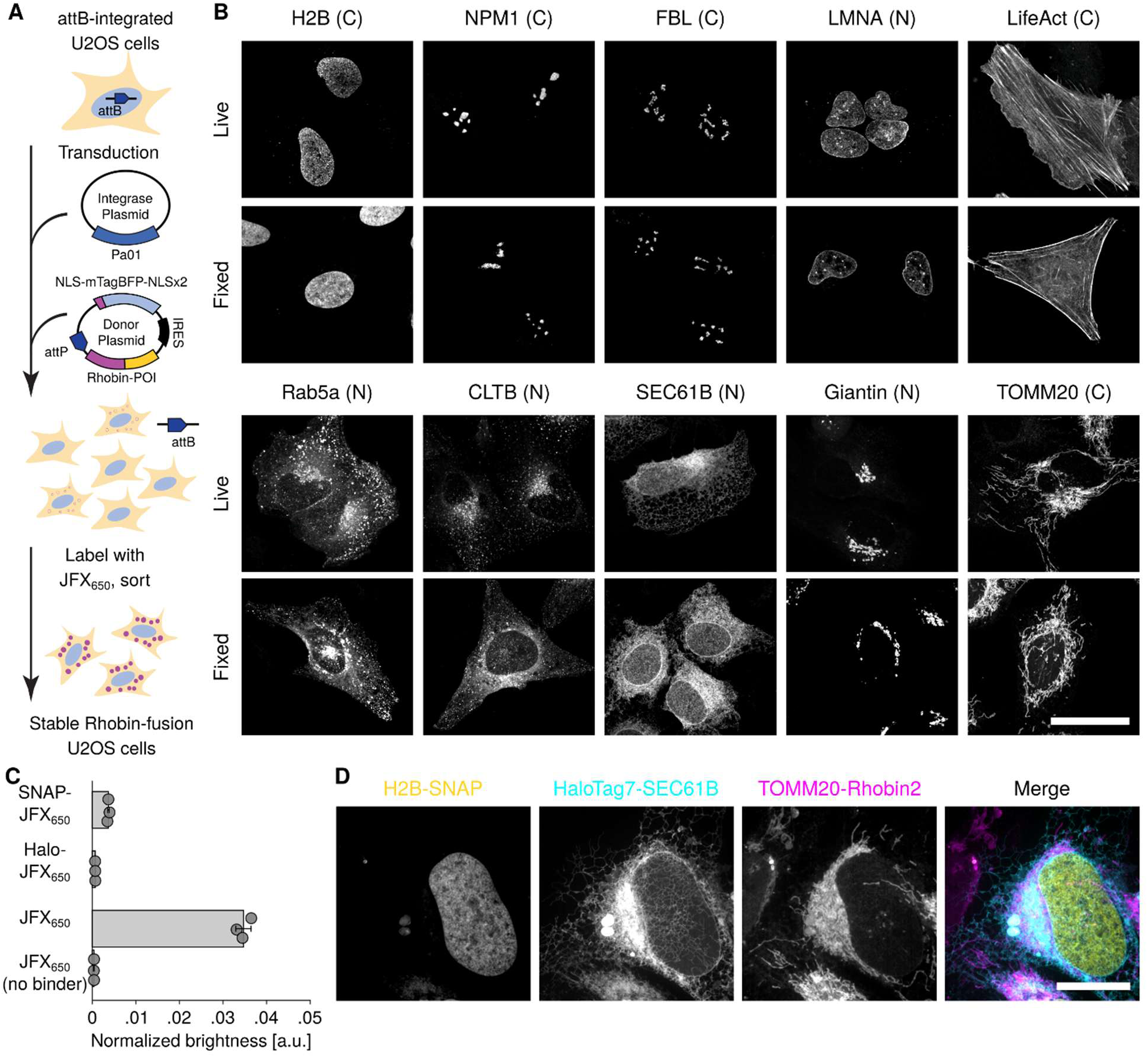
Multiplexed cellular imaging with Rhobin. (**A**) Schematic of U2OS stable cell line generation expressing Rhobin fusion proteins. Landing pad site attB-integrated U2OS cells were transduced with Pa01 integrase plasmid and donor plasmid encoding Rhobin fusion protein coding genes with attP sequence. After at least two passages, cells were labeled with JFX_650_ and sorted to isolate stable integrant cells. (**B**) Live-cell (top row) and fixed-cell (bottom row) SoRa spinning disk confocal imaging of Rhobin2-fusion proteins with 50 nM SiRh_p_. Protein fusion orientations are denoted as (C) for C-terminal and (N) for N-terminal tagging. (**C**) Quantitative analysis of Rhobin2 cross-reactivity with HaloTag ligand Halo-JFX_650_ and SNAP-tag ligand SNAP-JFX_650_. Individual data points and mean ± SD of normalized brightness of Rhobin2-H2B in cells labeled with JFX_650_, Halo-JFX_650,_ or SNAP-JFX_650_. See Table. S2 for numerical values. (**D**) Multiplexed live-cell SoRa spinning disk confocal imaging with triple labeling of nucleus (H2B-SNAP_f_ labeled with SNAP-JF_571_), ER (HaloTag7-SEC61B with Halo-JF_503_), and mitochondria (TOMM20-Rhobin2 with JFX_650_). All images are maximum intensity projections, scale bars: 20 μm, see table S3 for imaging conditions.

Rhobin2 also supports multiplexed imaging with other tags such as SNAP-tag and HaloTag. Rhobin2 prefers to bind JFX_650_ and it displayed lower intensity with JFX_650_-SNAP-tag ligand (∼10×) and JFX_650_-HaloTag ligand (∼50×; Fig. 3C). This allowed simultaneous multicolor imaging in cells expressing histone H2B-SNAP-tag (nucleus), HaloTag7-SEC61B (endoplasmic reticulum) and TOMM20-Rhobin2 (mitochondria) that were labeled with JF_571_ cpSNAP-tag ligand (*40*), JF_503_ HaloTag ligand (*20*), and Rhobin2 ligand JFX_650_ (Fig. 3D). These findings demonstrate the orthogonality of Rhobin and underscore its utility as a robust platform for both standalone and multiplex live-cell imaging.

### Advanced Fluorescence Microscopy Using Rhobin

The reversible dye binding by Rhobin enables exchange of photobleached fluorophores, which can compensate for the pronounced photobleaching that hampers photon-intensive live-cell imaging applications such as stimulated emission depletion (STED) microscopy (Fig. 4A). To test this, we incubated U2OS cells expressing LifeAct-Rhobin2 with JFX_650_ (100 nM) and performed both confocal and STED microscopy. The STED imaging experiments resolved individual actin fibers in actin bundles with higher resolution than confocal microscopy, and the exchangeable fluorophore system allowed imaging of actin dynamics over time (Fig. 4B-D, mov. S1). We also imaged fast ER dynamics using SEC61B-Rhobin2/SiRh_P_ as the label for over 100 time points and negligible photobleaching (Fig. 4E, mov. S2). Finally, we compared the performance of Rhobin with HaloTag7 or reHaloF with JFX_650_-HaloTag ligand, finding that the Rhobin labeling system showed the highest photostability (Fig. 4F-H).

**Fig. 4.**
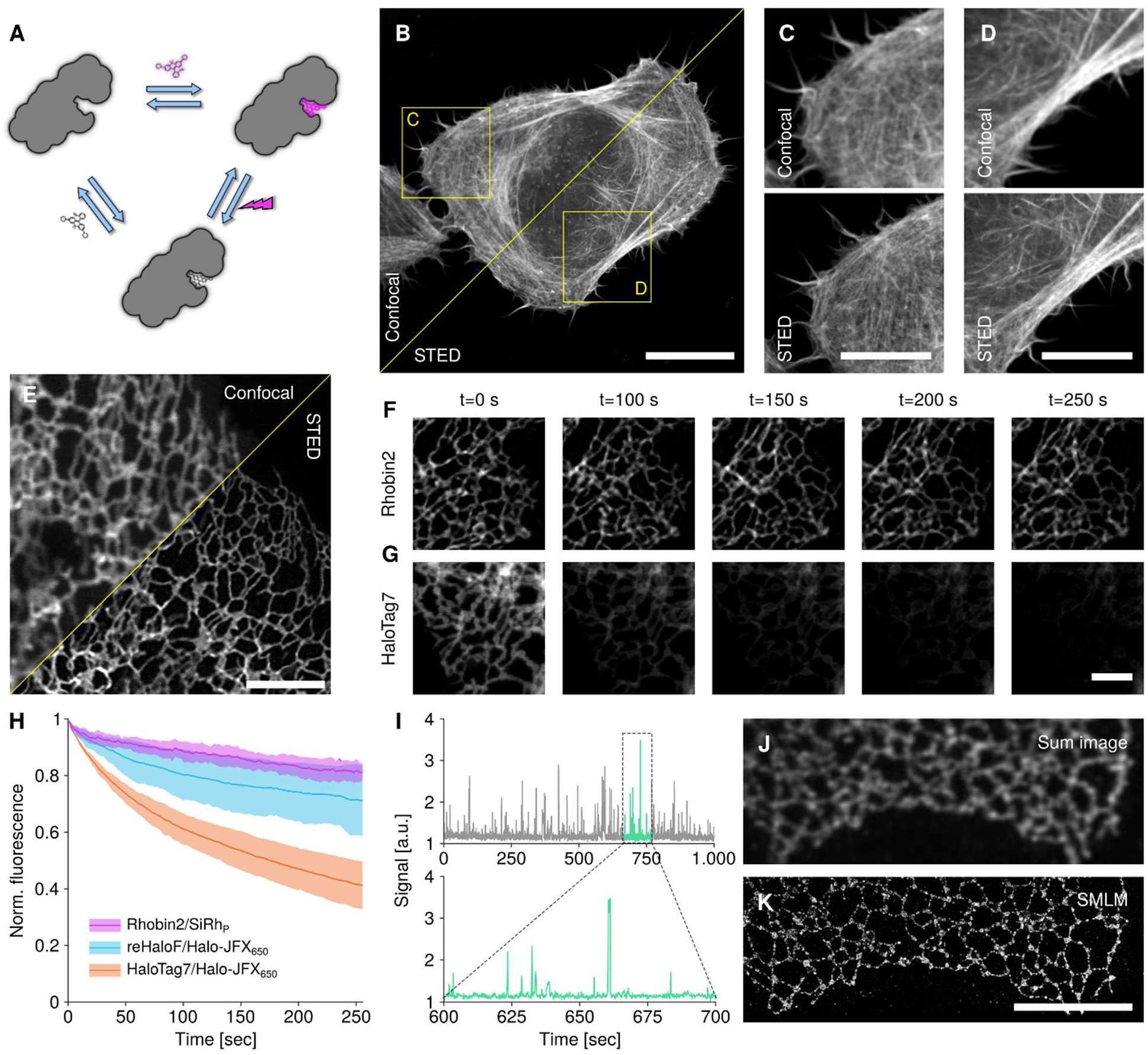
Super-resolution microscopy in live and fixed cells with Rhobin. (**A**) Rhobin transiently binds rhodamine molecules and thereby constantly replenishes signal lost through photobleaching of dyes. (**B-D**) No-wash STED microscopy of live U2OS cells transiently expressing LifeAct-Rhobin2 labeled with 100 nM JFX_650_. (C, D) Zoom-ins of regions highlighted in (B) imaged either with confocal or STED microscopy. Scale bar: 20 μm (B) and 10 μm (C, D). (**E-H**) Rhobin enables timelapse STED microscopy of fast ER dynamics with minimal signal loss. U2OS stably expressing N-terminal tag fusions of SEC61B were stained with 1 μM Halo-JFX_650_ (HaloTag7, reHaloF) or 2 μM SiRh_P_ (Rhobin2) and imaged at a frame rate of 0.387 fps for at least 100 time points. (F, G) Selected frames from timelapse acquisitions of Rhobin2:SEC61B labeled with SiRh_P_ (F) or HaloTag7:SEC61B labeled with Halo-JFX_650_ (G). Scale bar: 2 μm. (H) Signal loss during timelapse imaging. Mean ± SD signal inside cells over time for indicated constructs. (**I-K**) PAINT-type super-resolution microscopy with Rhobin. Rhobin2-SEC61B was transiently expressed in U2OS cells and labeled with 5 nM SiRh_P_ after chemical fixation. Imaging under no-wash conditions and near-TIRF illumination. (I) Repeated, but transient binding of SiRh_P_ molecules to Rhobin2 at low nanomolar concentrations can be observed as intensity bursts in intensity time traces extracted from a diffraction-limited area. See movie S3 for raw data of binding-induced blinking. (J) Sum image across 10,000 frames of an image stack, simulating a diffraction-limited image. (K). Reconstructed image from localized molecules in the full stack. Scale bar: 20 μm, see table S3 for imaging conditions.

The reversible binding of bright rhodamine dyes also facilitates points accumulation in nanoscale topography (PAINT) (*41*) experiments, a type of single-molecule localization microscopy (SMLM) (*42*). We labeled formaldehyde-fixed U2OS cells expressing SEC61B-Rhobin2 with 5 nM of the fluorogenic rhodamine SiRh_P_. At this low concentration, individual dye molecules transiently bound to Rhobin can be imaged via near total internal reflection fluorescence (TIRF) illumination and localized with sub-diffraction precision. The binding events appear as transient signals on the sub-second to second time scale over the entire duration of the measurement (Fig. 3I, mov. S3). Reconstruction of images from all localized binding events allowed us to obtain super-resolved images of the ER with improved resolution compared to widefield images (Fig. 4J,K).

### Protein-specific Labeling and Imaging in Extremophiles

We then tested the utility of Rhobin/dye pairs to label proteins in extremophile archaea. The harsh environment in which these organisms reside poses substantial challenges to labeling, resulting in few imaging studies (*43*, *44*). Consequently, the few currently available tagging approaches come with serious limitations and are not widely used. The different physiology of extremophiles makes them a desirable target for basic research, warranting the development of tools to study protein dynamics and subcellular structure in these cells. These organisms offer unique insights into the biochemical and molecular adaptations that enable life at the limits, shedding light on the evolutionary origins of complex cellular processes. Studying extremophiles also aids in the discovery of highly stable enzymes and biomolecules with broad industrial and biotechnological applications. The stability and binding of ligands at high temperatures make the Rhobin/dye system an attractive tag for this application.

To test how Rhobin performed under extreme conditions, we used *Sulfolobus acidocaldarius,* which requires culturing at 75 °C and pH 2 for growth, as the model system. We fused Rhobin9 to the cytoplasm-facing C-terminus of SlaB, one of the S-layer proteins in *S. acidocaldarius*, as well as to the N-terminus of the nucleoid-associated protein Cren7 (Fig. 5A). Expression of the fusion proteins did not affect cell growth (fig. S11). We imaged *S. acidocaldarius* expressing either SlaB-Rhobin9 or Rhobin9-Cren7 and stained with JF_660_ on a fluorescence microscope modified to maintain the sample at 75 °C (*45*). We achieved specific labeling of both fusion proteins with low background, with wild-type cells showing negligible nonspecific staining (Fig. 5B). SlaB-Rhobin9 exhibited a ring-like localization pattern in line with incorporation into the S-layer. Rhobin9-Cren7 is localized to a dense structure in the center of cells, consistent with incorporation into the nucleoid. This labeling system allowed following the dynamics of both Cren7 and SlaB during cell division through time-lapse imaging, enabling the observation of the timing of chromosome segregation and cell cleavage (Fig. 5C, movs. S4,5).

**Fig. 5.**
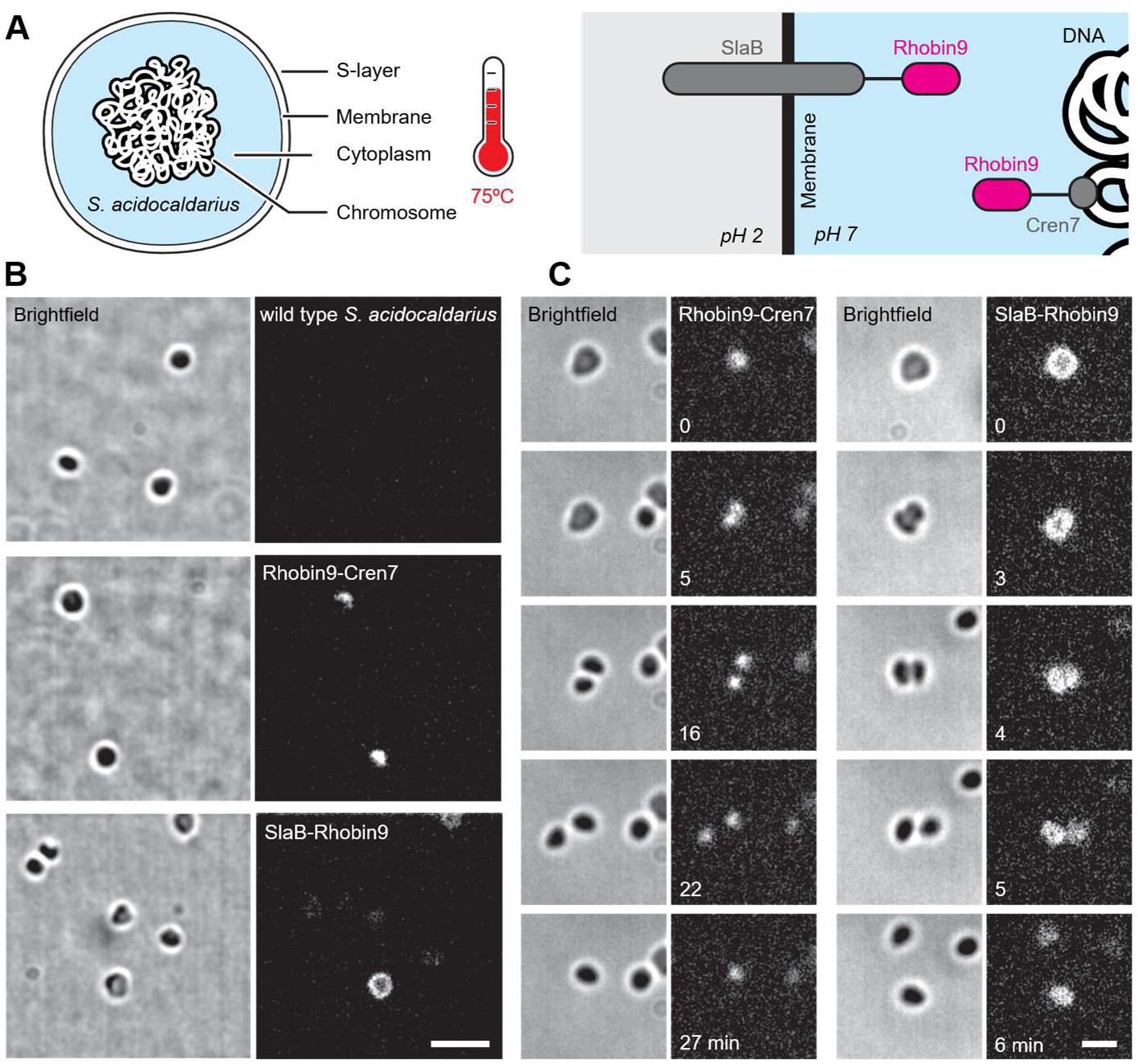
Rhobin enables protein-specific tagging in *S. acidocaldarus*. (**A**) Schematic of an archaea cell that lives at 75 °C and pH 2. Rhobin9 was genetically fused to either the S-layer transmembrane protein SlaB facing the cytosol or to the chromatin protein Cren7. (**B**) Transmitted light and fluorescence images of living *S. acidocaldarius* archaea cells, either wild type or expressing Rhobin9 fused to SlaB or Cren7. Samples were incubated with JF_660_, then washed with fresh media before imaging at 75 °C. (**C**) Timelapse imaging of dividing archaea cells. Chromosome segregation (Cren7) or cell cleavage (SlaB) can be observed (movies S4,5). Scale bars 1 μm, see table S3 for imaging conditions.

## Conclusion

Our design and application demonstration of Rhobin moves *de novo* protein design beyond the proof-of-concept stage. It is noteworthy that we designed Rhobin in a one-shot manner by testing only a handful of sequences without large-scale screening or directed evolution. We attribute this success to the understanding of the physical and chemical principles, the requirements of modern biological microscopy, and the use of advanced protein design methods. Looking forward, we expect Rhobin to be not just a useful tagging system for providing contrast in fluorescence microscopy, but also a basis for biosensor design.

## Acknowledgments

We thank all members of the DeGrado and Huang labs for helpful discussions. We thank the Gladstone Flow Cytometry Core for access to instrumentation and experimental support, the UCSF Innovation Core at the Weill Institute for Neurosciences for access to spinning disk, confocal, and STED microscopes, and the UCSF Center for Advanced Light Microscopy for access to TIRF microscopes. We thank the UCSF Small Molecule Discovery Center for access to their multimode plate reader, and Chad Altobelli for guidance on operating the plate reader. pSNAPf-H2B was a gift from New England Biolabs & Ana Egana (Addgene plasmid # 101124). pAF0033 Ef1a-Pa01-2a-EGFP was a gift from Patrick Hsu (Addgene plasmid #193460). Sniper2L was a gift from Hyongbum Kim (Addgene plasmid #193856). pET51b-His-TEV-HaloTag7 was a gift from Kai Johnsson (Addgene plasmid #167266). The pSVA-Ara-FX plasmid, MW001, and ER1821 cells were generous gifts from Sonja-Verena Albers (University of Freiburg, Germany).

## Funding

National Science Foundation (CHE-2108660 and MCB-2306190 to WDF)

National Institutes of Health (R35GM122603 to WDF, R01GM131641 to BH, and R35GM118119 to RDM)

Postdoc Fellowship of Cardiovascular Research Institute at UCSF to YC

Chan Zuckerberg Biohub San Francisco Investigator Program to BH

Howard Hughes Medical Institute to LDL and RDM

## Author contributions

Conceptualization: YC, KY, BH, WFD

Formal Analysis: YC, KY, KIH

Funding acquisition: YC, RDM, LDL, BH, WFD

Investigation: YC, KY, KIH, YK, AB, ACO, SJL

Methodology: YC, KY, KIH, YK, AB, ACO, SJL

Resources: YC, KY, KIH, YK, ACO, LDL, JBG

Software: YC, KY, LL, KL, KAH, SIM

Supervision: RDM, LDL, BH, WFD

Visualization: YC, KY, KIH, AB, SJL

Writing – original draft: YC, KY, KIH, BH, WFD

Writing – review & editing: YC, KY, KIH, AB, ACO, SJL, BH, WFD

## Data and materials availability

Plasmids used in this study are available from Addgene (accession IDs in Supplementary Materials) or upon request.

