## Supplementary Material for "*De novo* designed bright, hyperstable rhodamine binders for fluorescence microscopy"

### Materials and Methods

#### Computational Design

Generation of pan-rhodamine-specific scaffolds

To generate additional helical protein backbones beyond parametric helical bundles for binding rhodamines, we used a diffusion-based method to generate new protein scaffolds. Three inputs were used: 1) A conserved, well-packed protein base from previously designed four-helical bundles (*29*) (PDB: 7JRQ, residue 33-59 and 111-116) was used as input to bias the generation of helical structures; 2) An expanded rhodamine template with bulky substitutions was used to occupy the cavity space for diffused proteins, allowing subsequent application of such a cavity for binding rhodamines with a wide variety of minor substitutions. Its coordinate was modified from the rhodamine ligand (ID: PUJ) at PDB:6U2M using Pymol 3) To find the optimum residue for interacting with the carboxyl group present at the expanded rhodamine template and also almost all rhodamines, we picked Arg with a positive charge delocalized between two nitrogen atoms, providing an energetically-favorable solution to interact with carboxylic acids, carrying one negative charge delocalized between two oxygen atoms. The exact geometry of Arg relative to the expanded rhodamine template was guided by vdM database (*24*), where the Arg backbones with the lowest C score for interacting with the carboxyl group as a chemical group were used (*25*). The Arg-Rhodamine motif was placed at nine different positions at the extended core of a well-packed protein base, in close proximity to the original ligand position at PDB:7JRQ, which was optimized in the later step of partial diffusion.

With these starting poses, we ran RFdiffusion (*28*) to generate around 100 to 900 diffused backbones for each input. The example script for backbone generation was listed in Supplementary Text 1. Protein backbones were constructed using the expanded rhodamine template as the ligand to define the pocket space, a well-packed protein core, and the vdM-suggested Arg motif. The gaps between these motifs were bridged with connecting residues, resulting in proteins ranging from 131 to 171 residues in length.

The diffused backbones were first filtered by the steric clash with the expanded Rhodamine ligands superposed with the vdM geometry(*24*) (Supplementary Text 2). Next, the radius of gyration was calculated for each diffused backbone to select the ones with high compactness. Ligand SASA (solvent-accessible surface area) was calculated to select ones with sufficient ligand burial surface. The scaffolds passing these computational filters were evaluated manually with a preference for minimized fraction of nonbuttressed loops, high compatibility of protein pockets for binding rhodamines. pop The Rhobin scaffold (sequence length: 157 aa) was selected based on all these computational filters and principal considerations.

Partial diffusion (*28*) (Supplementary Text 3) was used to generate 2000 similar scaffolds with the selected Rhobin scaffold to improve the designability of Rhobin scaffolds and enlarge the local backbone diversity of the Rhobin scaffolds. The 49 Rhobin-similar scaffolds with minimal clash with the vdM-placed expanded rhodamine template were used as scaffolds for sequence design.

Sequence design for Rhobins binding to $JF_{660}$

To design binders for $JF_{660}$, we first placed $JF_{660}$ into the 49 Rhobin-similar scaffolds based vdM geometry (Supplementary Text 2), which is the carboxyl-Arg geometry we used previously at scaffold generation. To improve protein stability, we removed the 6 residues from the N-terminal and 7 residues from the C-terminal, because they are less packed than the other residues in the core, which also minimized the sequence to 144 aa. Sequence design was carried out with 3 iterative rounds of LigandMPNN and Rosetta FastRelax for 1000 sequences per input (*26*, *46*). Constraints were applied to Rosetta FastRelax to maintain the low-energy conformation of $JF_{660}$ (Supplementary Text 4). We ran the sequence design with fixation of Arg residue at residue 80, affording the original vdM used for RFdiffusion at residue 80. To access a broad sequence space to interact with the carboxylic group of $JF_{660}$, we also did the sequence design without the amino acid constraint at residue 80. exploring the potential of other interaction residues. All the designed sequences were subjected to structure prediction with RaptorX-Single.(*33*) The selection criteria were the sub-Å backbone RMSD between the designed complex and the predicted unliganded structures, and also agreement of sidechain conformation of active site between design and prediction, High pLDTT (>85), and low backbone RMSD (< 1 Å). Potential hydrogen bonding to the carboxylic acid of the $JF_{660}$ was also used as a filter to rule out the designs without polar atoms around 3.5Å of the carboxylic group of $JF_{660}$ (Supplementary Text 5). Designing passing these computational filters were evaluated manually for 1) maximal saturation of hydrogen bond potential of carboxyl group of $JF_{660}$ with two or more polar interactions, 2) minimal hydrophobic packing around the carboxylic acid group of $JF_{660}$ to disfavor its lactonization, because free rhodamines of lactone form were favored in solvents of low dielectric point (*20*). 3) inclusion of second-shell interactions to stabilize the rotamer of the first-shell interacting residues. None of the R80-containing sequences were selected for experimental validation, based on these criteria and the potential bias of the selected Rhobin scaffold.

**Bacterial expression and *in vitro* characterization**

Bacterial expression of Rhobin

The plasmids encoding Rhobin, cloned into pET29b(+) vector with NdeI and XhoI restriction sites, were transformed into *E. coli* BL21(DE3) (C2527H, New England Biolabs) and plated on an LB agar plate with 50 µg $ml^{-1}$ kanamycin (Thermo Scientific). This same kanamycin concentration was used throughout the experiment. Single colonies were inoculated into 5 mL LB medium containing kanamycin and grown at 37 °C, 220 rpm for 5-6 hours. 5 ml of starter culture was then diluted into 0.2 L of LB with kanamycin and allowed to grow at 37 °C until $A_{600}$ reached 0.6-0.8. The culture was induced upon the addition of 0.5 mM isopropyl-ß-D-1-thiogalactopyranoside (IPTG) (Apex Bioresearch Products) and grown at 30 °C for 20 hours. Cells were harvested by centrifugation (Sorvall Legend RT+, Thermo Scientific) at 4 °C, 4, 3500 rpm for 15 min., flash frozen in liquid nitrogen, and stored at -80 °C.

For protein extraction, the cell pellets were resuspended in PBS buffer (119-069-13, Quality Biological) with added imidazole for a final concentration of 20 mM and lysed via sonication. The resulting cell lysates were clarified by centrifugation at 14,000 g for 30 min. The supernatants after centrifugation were purified with nickel-nitrilotriacetic acid (Ni-NTA) resin (Invitrogen), following the manufacturer's instructions. After buffer exchange to PBS using a PD-10 column (17085101, Cytiva), proteins were analyzed using sodium dodecyl-sulfate polyacrylamide gel electrophoresis (SDS-PAGE) and fast protein liquid chromatography (FPLC).

Bacterial expression of HaloTag7 was conducted with pET51b-His-TEV-HaloTag7 (Plasmid #167266) purchased from Addgene according to the reported protocol (*47*).

Binding constant determination of Rhobin with $JF_{660}$

We used fluorescence titration experiments to determine the binding dissociation constants of Rhobin for $JF_{660}$ with BioTek Synergy Neo2 Reader. $JF_{660}$ binding to Rhobin caused a fluorescence increase (Ex: 650/20 nm, Em: 691/20 nm) in 8 of 9 designed Rhobin, which was used to quantify the bound $JF_{660}$. For each Rhobin, protein in PBS was serially diluted while fixing the concentration of $JF_{660}$ at 0.2 μM. The single-site binding model (Equation 1, $N = 1$, $L = 0.2$) was used to fit $K_D$ using GraphPad Prism, considering the total fluorescence from both bound $JF_{660}$ and free $JF_{660}$ (Equation 2). $b_{free}$ was obtained by measuring the fluorescence of the $JF_{660}$ in PBS after serial dilution.

***(Equation 1)***

$$\text{bound ligand} = \frac{1}{2}\left[K_D + [Ligand_{\text{total}}] + \frac{[Protein_{\text{total}}]}{N} - \sqrt{\left(K_D + [Ligand_{\text{total}}] + \frac{[Protein_{\text{total}}]}{N}\right)^2 - 4\,[Ligand_{\text{total}}]\frac{[Protein_{\text{total}}]}{N}}\,\right]$$

- $Protein_{total}$: total protein concentration
- N: binding stoichiometry
- $K_D$: binding constant of protein-ligand interaction

***(Equation 2)***

$$Fluorescence = b_{bound}[Ligand_{bound}] + b_{free}[Ligand_{free}]$$

- $Ligand_{total:}$ total concentration of ligand
- $Ligand_{free:}$ free ligand
- $Ligand_{bound}$: ligand binding to protein
- $Ligand_{free} = Ligand_{total} - Ligand_{bound}$
- $b_{free}$ = brightness (fluorescence) of free ligand per μM
- $b_{bound}$ = brightness (fluorescence) of bound ligand per μM

Thermal stability

For protein characterization assays, proteins were prepared in 10 μM concentrations in PBS buffer. CD spectra were collected on a Jasco J-810 CD spectrometer in a 0.1 cm path-length quartz cuvette. The parameters for full spectra collections were set up as follows: the bandwidth was set to 2 nm, the scanning speed was set to 50 nm/min, and the average of accumulations was set to 3. Temperature-dependent data were collected at 222 nm from 20 to 95 °C with an interval of 5 °C and an increase rate of 2 °C/minute.

For ligand binding assays, 20 μM Rhobin9 was incubated with 10 μM $JF_{660}$ at room temperature for 10 minutes before transferring to a 1 cm path-length quartz cuvette. As the control, 20 μM HaloTag7 was incubated with 10 μM $JF_{660}$-HaloTag7 ligand (synthesized by Luke Lavis lab) at 37 degree for 30 minutes to facilitate the covalent bond formation (https://pubs.acs.org/doi/10.1021/acs.biochem.1c00258) CD spectra for $JF_{660}$ ( 630 nm to 690 nm) in the form of noncovalent complex with Rhobin9 and covalent complex with HaloTag7 were collected firstly at 20 degree on a Jasco J-810 CD spectrometer. Next, the temperature was raised to 75 degrees with a waiting time of 2 minutes to collect the spectra with the same collection parameters. Collection parameters are as follows: the bandwidth was set to 4 nm, the scanning speed was set to 20 nm/min, the response time was set to 8 s, the pitch was set to 1 nm, and the average of accumulations was set to 5.

**Experiments in mammalian cells**

Cloning of mammalian expression plasmids

All plasmids encoding Rhobin or other fusion proteins used for the transient overexpression in mammalian cells were cloned into a nuclear marker co-expressing a custom backbone vector, generated by the insertion of the IRES-NLS-mTagBFP-NLSx2 gene fragment into pCMV OriP (Twist Bioscience) with In-Fusion Snap assembly master mix (TaKaRa). Rhobin coding genes were amplified by PCR from plasmids for bacterial expression, and coding genes for other fusion proteins, such as other fluorescent tags/proteins or subcellular localization domains, were assembled into the backbone vector with In-Fusion Snap assembly master mix. For cloning landing pad donor plasmids, the Pa01 attP site was inserted by overlapping PCR and assembled by In-Fusion cloning with In-Fusion Snap assembly master mix.

Cloned plasmids were transformed into HST08 Stellar competent *E. coli* (TaKaRa), and purified plasmids were verified by whole plasmid sequencing (Plasmidsaurus).

Mammalian U2OS cell culture

U2OS wild-type (ATCC) or genome-edited U2OS stable cells were cultured in either CELLSTAR T-25 flasks (Greiner Bio One) or 6-well plates (Falcon) with McCoy's 5A modified medium (Gibco) supplemented with 10% FBS (UCSF Cell Culture Facility), 1x GlutaMax (Gibco), 1 mM Sodium Pyruvate (HyClone), and Streptomycin/Penicillin (UCSF Cell Culture Facility) in a 37°C and 5% $CO_2$ humidified incubator.

For transient transfection, U2OS cells were seeded in 8-well chambered coverglasses or 96-well glass-bottom plates, then incubated at 37 ºC, 5% $CO_2$ in a humidified incubator (Panasonic) for 12-24 hours. 200 ng of plasmid DNA was transfected with JetOptimus (Polyplus) as per the manufacturer's instructions, then the media was replaced 4 hours after transfection. Transfected cells were incubated in a 37 ºC, 5% $CO_2$ humidified incubator for 12-24 hours prior to imaging.

Stable cell line generation

U2OS landing pad (LP) cell lines harboring Pa01 attB at the *AAVS1* safe harbor locus were generated as described previously (*48*). To integrate coding genes into landing pad cell lines, a donor plasmid containing sequences of Pa01 attP-Rhobin-fusion protein and an integrase-expression plasmid pEf1a-Pa01 were co-transfected into U2OS LP cells. For this, 250,000 U2OS LP cells were seeded on 6-well plates (Falcon), then incubated at 37 ºC, 5% $CO_2$ until reaching ~ 90% confluency. 1,000 ng of each donor and integrase-expressing plasmids were co-transfected with JetOptimus as per the manufacturer's instructions. Media was replaced 4 hours after transfection and cells were incubated at 37 ºC, 5% $CO_2$. After 2 days of incubation, cells were trypsinized with 0.25% Trypsin-EDTA (UCSF Cell Culture Facility), then passaged with fresh media on a new 6-well plate. After at least two further passages, genome-edited cells were trypsinized, washed with PBS (Gibco), then resuspended in collection buffer (10% FBS, 1x Streptomycin/Penicillin in PBS) supplemented with 200 nM $JFX_{650}$ or $SiRh_P$ and incubated for at least 20 minutes. $JFX_{650}$-positive cells were enriched by fluorescence-activated cell sorting (BD

FACS Aria Fusion 1) and collected in collection buffer (10% FBS, 1x Streptomycin/Penicillin in PBS). Sorting was validated by widefield fluorescence microscopy.

Sample preparation for live- and fixed-cell imaging

Cultured U2OS Rhobin-fusion protein stable cells or U2OS wild-type cells were trypsinized and resuspended in McCoy's 5A modified medium. 8-well chambered coverglasses (Lab-Tek II, Nunc) or glass-bottom 96-well plates (Cellvis) were passivated with poly-L-lysine (Sigma-Aldrich) for 30 minutes, then washed twice with PBS before cell seeding. Cells were seeded at a density of 20-50,000 cells/well and grown for 12-24 hours prior to transient transfection or imaging. Before imaging, the media was discarded, and cells were washed twice with PBS. Media was further changed to imaging media (FluoroBrite DMEM supplemented with 10% FBS and 1x GlutaMax, all Gibco or UCSF Cell Culture Facility), and samples were stained for at least 1 hour at 37°C, 5% in a humidified incubator before imaging. Samples were imaged under no-wash conditions unless noted otherwise.

For the preparation of fixed cells, the media was discarded, and cells were washed twice with DPBS (UCSF Cell Culture Facility). Cells were fixed with 3.2% PFA in DPBS at RT for 15 minutes. After fixation, cells were washed twice with DPBS, then labeled with rhodamine dye in DPBS solution for an hour at RT before imaging.

Triple labeling for multiplexed live-cell imaging

25,000 of U2OS TOMM20-Rhobin2 stable cells were seeded in a poly-L-lysine-passivated 8-well chambered cover glass 12-24 hours before imaging. 100 ng each of pSNAPf-H2B and pHaloTag7-SEC61B-IRES-NLS-mTagBFP-NLSx2 were transfected with JetOptimus as per the manufacturer's instructions, then the media was replaced 2 hours post-transfection. 8 hours post transfection, H2B-SNAPf and HaloTag7-SEC61B were covalently labeled with 100 nM Halo-$JF_{503}$ and 1 μM SNAP-$JF_{571}$ in imaging media. After 1 hour of labeling, cells were washed twice with imaging media, then labeled with imaging media with 100 nM $JFX_{650}$ for an hour and imaged under no-wash conditions

Widefield fluorescence microscopy

For widefield fluorescence microscopy, samples were prepared as described above. Imaging was performed on a custom-built microscope constructed around a Ti-E body (Nikon) equipped with a motorized stage (MS-2000, ASI), an active focus stabilization system (PFS, Nikon), a water immersion objective (CFI Plan Apochromat IR 60x WI NA 1.27, Nikon) and a LED light source (X-Cite XLED1, Excelitas). A sCMOS camera (Orca Flash 4.0, Hamamatsu) with a back-projected pixel size of 108 nm was used for acquiring images. Emitted fluorescence was separated from excitation light using a 405/488/561/640 nm quadband dichroic. 450/50, 525/50 and 700/75 nm bandpass filters were used to detect signal in the mTagBFP, mNeonGreen and far-red channel respectively. frFAST with its ligand HPAR-3OM was excited at 561 nm, and the emitted signal was filtered with a 700/75 nm bandpass filter. Samples were maintained at 37 °C and 5% $CO_2$ using a stage-top incubation chamber with an environmental control unit (Tokai Hit). All components of the imaging setup were controlled using the MicroManager 2.0 software platform and custom-written acquisition scripts (*49*).

Data analysis for widefield screening microscopy

Multi-channel image stacks were processed with a custom-written framework implemented in MATLAB 2024b. First, nuclei in individual multi-channel image stacks were segmented using the Matlab-Cellpose v2.0 bridge (*50*) using the mTagBFP channel as input. Segmented nuclei were filtered according to size, removing the 20% smallest and largest regions of interest (ROIs) to remove instances where segmentation led to incomplete segmentation of nuclei or erroneous merging of multiple nuclei into a single ROI. Next, the mean intensities in the mTagBFP, mNeonGreen, and far-red (protein tag ligands) channels were extracted for each mask. Intensities were offset-corrected, using pre-determined offsets for each channel determined in an empty sample not showing any specific fluorescence signal. Scatter plots of per-cell intensities in the mTagBFP vs mNeonGreen channels were used to measure expression level. Scatter plots of per-cell intensities in the mNeonGreen vs far-red fluorophore signal were used to measure the expression level normalized far-red fluorophore signal generated by fluorophores attaching to protein tags in the nucleus. Cells from each condition were subjected to linear regression model fitting using the MATLAB function 'fitlm'. Typical analyses contain >1,000 cells per condition.

SoRa spinning-disk confocal microscopy

Images were acquired using a CSU-W1 SoRa spinning disk confocal microscope (Weill Institute for Neuroscience, UCSF), constructed around a DMi8 inverted microscope body (Leica) equipped with a CSU-W1 SoRa confocal scanner unit (Yokogawa, 50 µm pinhole, SoRa disk with uniformizer), a motorized stage with Piezo top plate (ASI), a Plan Apo 63x/NA 1.40 oil immersion objective lens (Leica), a Kinetix sCMOS camera (Photometrics) and 401, 487, 561, 639 nm excitation lasers (Vortran). Samples were mounted in a stage-top 37 °C, 5% $CO_2$ incubation chamber with an environmental control unit (Okolab). The microscope was operated using Micro-Manager 2.0 software. Typically, z-stacks with 31 slices and 0.2 µm step size were recorded in 63x SoRa mode (pixel size in sample space: 26.6 nm). Maximum projections were generated with ImageJ. For subcellular imaging, each image was acquired with 200 ms exposure time per slice and 639 nm excitation at 4 mW. For multiplex imaging, 3-channel images with 200 ms exposure time per slice/channel were acquired at 488, 561, and 639 nm excitation with 1.5, 1,5, and 4 mW excitation power, respectively.

Confocal and STED microscopy

STED and point-scanning confocal images were acquired on a Stellaris8 TauSTED microscope (Leica) equipped with an 86x 1.2NA water immersion and a 100x 1.4NA oil immersion objective, HyD-X/R detectors, a pulsed white light laser (WLL) for excitation, and a 775 nm STED depletion laser. Samples were maintained at 37 °C and 5% $CO_2$ using an environmental control enclosure during imaging. Samples were excited with the WLL at 638 nm with a 638 nm notch filter added to the excitation light path, and the WLL was operated at a base power output of 85%. The emitted signal was collected with the HyD-R detector operating in photon counting mode and recording signal in the 642-740 nm spectral window with a pinhole size of 1 airy unit. tauSTED gating was applied for all STED acquisitions. Samples were scanned in unidirectional scanning mode with variable scan speeds and line averaging (see below).

STED images were acquired with two different settings: Slower acquisition for a larger field of view (Figs. 4B,D, mov. S1) and fast acquisition for a smaller field of view (Figs. 4E-G, mov. S2). Slow STED timelapse images with a size of 2048x2048 pixels were acquired using the 86x water immersion objective at a pixel size of 40 nm and a pixel dwell time of 20 µs with 2D STED depletion at 1.208 W nominal output power. No line or frame averaging was used. Fast STED timelapse images with a size of 512×512 pixels were acquired using the 100× oil immersion objective at a pixel size of 40 nm and a pixel dwell time of 2.8 µs with 2D STED depletion at 0.338 W nominal output power. 2× line averaging was applied.

Confocal images were acquired with identical settings, but without STED depletion and at slightly reduced excitation intensities to avoid detector saturation.

Single-molecule localization microscopy

Samples for single-molecule localization microscopy were prepared as described above. U2OS cells, transiently expressing Rhobin2-SEC61B were chemically fixed and labeled with 5 nM SiRhP for at least 1 hour.

Imaging was performed on a N-STORM TIRF microscope constructed around a Ti-E stand (Nikon) and equipped with an Apo TIRF 100×/1.49NA objective (Nikon), active focus stabilization (perfect focus system, Nikon), an iXon 897 EMCCD camera (Andor), and a LUD-H series laser unit with 405/488/561/647 nm excitation lasers (Nikon). The emitted light was filtered through an ET705/72m bandpass filter (Chroma). Prior to recording data, the illumination lasers were centered onto the optical axis of the objective, and the coupling angle was adjusted to achieve near-TIRF illumination. For each acquisition, 10.000 images were recorded with an exposure time of 100 ms at minimum interval, maximum laser power at 647 nm, and the electron multiplying gain set to 500.

Raw data was processed with the ImageJ plugin ThunderSTORM (*51*) using Wavelet image pre-filtering and maximum likelihood fitting of 2D emitter positions without multi-emitter fitting. Reconstructed images were rendered using the average shifted histogram approach with a final pixel size of 10 nm.

**Experiments in *Sulfolobus acidocaldarius***

Expression of Rhobin9 fusion protein in *Sulfolobus acidocaldarius*

Rhobin9 was fused to the C-terminal, cytoplasmic end of the S-layer protein SlaB or to the Nterminus of the nucleoid-associated protein Cren7 from *S. acidocaldarius*. In both cases, a xylose-inducible promoter *P*xyl was included upstream of the start codon, and the linker GGTGGGGSGG was codon optimized for *Sulfolobus acidocaldarius* and inserted between Rhobin9 and the target protein. Constructs were ligated between the SacII and NotI sites of the pSVAara-FX vector (*52*), yielding pACO48 (SlaB-linker-Rhobin9) and pACO49 (Rhobin9linker-Cren7). Plasmids were methylated via passaging in *E. coli* ER1821 (New England Biolabs, Frankfurt am Main, Germany) and transformed in electrocompetent *Sulfolobus acidocaldarius* MW001 cells. (*53*). Protein expression was induced by the addition of 0.2% D-xylose.

Cell culture and preparation of conditioned media

The *S. acidocaldarius* strain MW001 (wild type) was grown aerobically in Brock's minimal media (pH 3) (*54*) supplemented with 0.1% tryptone (Sigma-Aldrich), 0.2% sucrose, and 10 µg/mL uracil. For strains carrying plasmids, uracil was omitted from the medium, while all other conditions remained the same. Cultures were incubated at 75 °C with shaking, and growth was monitored by measuring optical density at 600 nm ($OD_{600}$). To prepare conditioned medium, 50 mL of an exponentially growing culture ($OD_{600}$ ~0.3–0.4) was centrifuged at 4,000 × g for 15 minutes at room temperature, and the resulting supernatant was filtered through a 0.22 µm filter to remove any remaining cells or debris. The filtered conditioned medium was stored at 4 °C for up to one month.

Cell growth and labeling

50 ml cultures of *S. acidocaldarius* MW001 strains harboring the plasmids pACO48 (*slaB-linker-rhobin9*), pACO49 (*rhobin9–linker-cren7*), or no plasmid (WT) were grown overnight at 76 °C in an incubator with shaking until an OD of 0.3–0.4 was reached. The cultures were then induced with 0.2% xylose for 4 h to express the Rhobin9-tagged protein constructs. 1 ml aliquots of the cultures were collected in microfuge tubes, supplemented with 250 nM $JF_{660}$ dye, and incubated at 76 °C in a benchtop shaker (300 rpm) for 10 minutes. The cells were then pelleted by centrifugation at 4,000 g for 3 minutes and washed twice with Brock's medium containing 0.2% xylose. After washing, the pellets were resuspended in 1 mL of filtered conditioned medium and incubated at 76 °C with shaking (300 rpm) for 20 minutes to recover.

Sample preparation for imaging

Samples for imaging were prepared following a previous protocol (*45*). To promote cell adhesion, an equal volume of freshly boiled 1.7% Gelrite (Gelzan, Sigma) was combined with 2× Brock's minimal medium preheated to 75 °C. This mixture was used to coat the bottom surface of a Delta T imaging chamber by manual streaking with a sterile pipette tip. After allowing the coating to solidify for 5 minutes at room temperature, 2 mL of conditioned Brock's medium was added to the chamber. Prepared chambers were kept in an incubator at 75 °C until used for imaging. Prior to imaging, the microscope objective was preheated to 70 °C and kept at this temperature throughout the experiment. An aliquot of 500 µL of recovered culture was transferred to the prepared chamber (final volume ~2.5 mL) and placed on the microscope stage immediately to minimize thermal fluctuations. The chamber temperature was maintained between 75-76 °C using a temperature controller set to 76 °C. The same controller was used to adjust the temperature of the glass lid until condensation was no longer observed.

iSIM Imaging of *S. acidocaldarius*

Images were acquired using Micro-Manager on a Nikon Ti-E microscope with Perfect Focus, an ASI MS-2000 XY piezo Z stage, a Hamamatsu Quest camera, and VT-iSIM with Ingwaz and 100200mW 405, 488, 561, 642 nm lasers and a Cairn Optospin emission filter wheel with ET 450/50m, ET 525/50m, ET 595/50m, ET 655lp, and ZET 405/488/651/640m Chroma filters. Live cells were imaged using a Bioptechs DeltaT- incubator set to 76 °C and an objective heater set to 70 °C on a Nikon 100x 1.40NA PlanApo objective, as described previously (*45*).

**Supplementary text 1: Example of RFdiffusion job file for generating Rhobin scaffolds**

T../../scripts/run_inference.py
inference.output_prefix=example_outputs_900_1/Suffix
inference.input_pdb=input/diffusion_input.pdb
'contigmap.contigs=[20-30/A33-59/25-35/B200-202/10-20/A91-116/20-30]'
inference.num_designs=800
potentials.guide_scale=5
'potentials.guiding_potentials=["type:substrate_contacts,s:1,r_0:8,rep_r_0:5.0,rep_s:2,rep_r_min:1"]' potentials.substrate=jf6 inference.ckpt_override_path=../../models/ActiveSite_ckpt.pt

**Supplementary text 2: Placing ligand with vdM geometry.**

```
from Bio import PDB
import os
def find_consecutive_drf_atoms(structure):
        """Finds consecutive residues D-R-F and returns their Cα atoms."""
        ca_atoms = []
        residues = list(structure.get_residues())
        for i in range(len(residues) - 2):
            res1, res2, res3 = residues[i], residues[i+1], residues[i+2]
            # Check if the residue names match 'D', 'R', 'F'
            if res1.get_resname() == 'ASP' and res2.get_resname() == 'ARG' and res3.get_resname()
== 'PHE':
                        try:
                                    ca_atoms = [res1["CA"], res2["CA"], res3["CA"]]
                                    return ca_atoms  # Return first match
                        except KeyError:
                                    # If any CA atom is missing, skip
                                    pass
            return None  # If no matching DRF sequence found

        def get_ca_atoms_by_resnums(structure, residue_numbers):
            """Extract Cα atoms from given residue numbers."""
            atoms = []
            for residue in structure.get_residues():
                if residue.id[1] in residue_numbers:
                    try:
                        ca_atom = residue["CA"]
                        atoms.append(ca_atom)
                    except KeyError:
                        pass
            return atoms

        def extract_residue(structure, chain_id, residue_id):
            """Extract a specified residue by chain and residue number."""
            for chain in structure.get_chains():
                if chain.id == chain_id:
                    for residue in chain:
                        if residue.id[1] == residue_id:
                            return residue
            return None

        def add_residue_to_structure(structure, residue, chain_id):
            """Add a residue to a specified chain."""
            new_residue = residue.copy()
```

```
    model = structure[0]  # Assuming first model
    if chain_id in model.child_dict:
        chain = model[chain_id]
    else:
        chain = PDB.Chain.Chain(chain_id)
        model.add(chain)

    residue_id = new_residue.id
    if residue_id in chain.child_dict:
        print(f"Warning: Residue ID {residue_id} already exists in chain {chain_id}.")
    else:
        chain.add(new_residue)

# === Main settings ===
reference_pdb_path =
'/wynton/home/degradolab/yuda/RFdiffusion/demo/diffusion7a/input/diffusion7a.pdb'
target_pdb_directory =
'/wynton/home/degradolab/yuda/RFdiffusion/demo/diffusion7a/example_outputs_900_1'
output_directory =
'/wynton/home/degradolab/yuda/RFdiffusion/demo/diffusion7a/example_outputs_900_1_jf6'

# Create output directory
if not os.path.exists(output_directory):
    os.makedirs(output_directory)

# Load reference structure
parser = PDB.PDBParser()
ref_structure = parser.get_structure('reference', reference_pdb_path)

# Extract CA atoms from residues 85-87 of reference
ref_residue_numbers = [200, 201, 202]
ref_ca_atoms = get_ca_atoms_by_resnums(ref_structure, ref_residue_numbers)

# Extract ligand (residue 1 in chain X)
ref_residue = extract_residue(ref_structure, 'X', 1)
if ref_residue is None:
    raise ValueError("Specified residue not found in reference structure.")

# Process target PDBs
for pdb_file in os.listdir(target_pdb_directory):
    if pdb_file.endswith(".pdb"):
        target_path = os.path.join(target_pdb_directory, pdb_file)
        target_structure = parser.get_structure('target', target_path)

        # Find DRF sequence in target structure
```

```
target_ca_atoms = find_consecutive_drf_atoms(target_structure)

if target_ca_atoms is not None and len(target_ca_atoms) == len(ref_ca_atoms):
    # Superimpose based on DRF <--> 85-87
    super_imposer = PDB.Superimposer()
    super_imposer.set_atoms(ref_ca_atoms, target_ca_atoms)
    super_imposer.apply(target_structure.get_atoms())  # Apply transformation

    # Add ligand from reference to target
    add_residue_to_structure(target_structure, ref_residue, 'X')

    rmsd = super_imposer.rms
    print(f"RMSD for {pdb_file}: {rmsd:.3f}")

    # Save aligned structure
    io = PDB.PDBIO()
    io.set_structure(target_structure)
    io.save(os.path.join(output_directory, pdb_file))
else:
    print(f"Warning: DRF sequence not found or incomplete in {pdb_file}")
```

**Supplementary text 3: Partial diffusion of the selected Rhobin scaffold**

```
../../scripts/run_inference.py
inference.output_prefix=output_2000/suffix
inference.input_pdb=input/protein.pdb
'contigmap.contigs=[61/A62-63/21/A85-87/10/A98-99/34/A134-135/22]'
inference.num_designs=2000
diffuser.partial_T=25
```

**Supplementary text 4: Constrain for Rosetta FastRelax**

```
Dihedral C8 1X C7 1X C4 1X C3 1X CIRCULARHARMONIC 1.5 0.4
```

### Supplementary text 5: Potential HB around the carboxylic group of JF660

```
import os
import prody as pr
import csv

# Define the directory containing the PDB files
pdb_dir =
'/wynton/home/degradolab/yuda/LigandMPNN/ligandMPNN_rosettaFR_pipeline/rhobinder_2e3_202405/method1_
vdm/conf_a/output/final_out/'

# Specify the ligand residue name and the atoms of interest
ligand_resname = '60a'  # Replace 'LIG' with your ligand residue name
specified_atoms = ['O1', 'O2']  # Replace with the actual atom names in your ligand

# Specify the output CSV file
output_csv =
'/wynton/home/degradolab/yuda/LigandMPNN/ligandMPNN_rosettaFR_pipeline/rhobinder_2e3_202405/method1_
vdm/conf_a/analysis/atoms_around_coo_60a.csv'

# Function to process each PDB file and return atom counts and residue details
def process_pdb_file(pdb_path):
    pdb = pr.parsePDB(pdb_path)
    ligand = pdb.select(f'resname {ligand_resname}')
    if not ligand:
        return None

            ligand_atoms_of_interest = ligand.select('name ' + ' '.join(specified_atoms))
            if not ligand_atoms_of_interest:
                return None

            atom_counts = {'C': 0, 'N': 0, 'O': 0}
            residue_details = {'Carbon': [], 'Nitrogen': [], 'Oxygen': []}

            for atom in ligand_atoms_of_interest:
                neighbors = pdb.select(f'within 3.5 of index {atom.getIndex()} and protein and (element C or element N
or element O)')
                if neighbors:
                    for neighbor in neighbors:
                        atom_type = neighbor.getElement()
                        residue_number = neighbor.getResnum()
                        residue_name = neighbor.getResname()

                        if atom_type in atom_counts:
                            atom_counts[atom_type] += 1
                            if atom_type == 'C':
                                residue_details['Carbon'].append((residue_number, residue_name))
                            elif atom_type == 'N':
                                residue_details['Nitrogen'].append((residue_number, residue_name))
                            elif atom_type == 'O':
                                residue_details['Oxygen'].append((residue_number, residue_name))
                            else:
                                # Handle unexpected atom types here if needed
                                pass

            return atom_counts, residue_details
```

```
    # Prepare to write to CSV
    with open(output_csv, mode='w', newline='') as file:
        writer = csv.writer(file)
        writer.writerow(['PDB File', 'Carbon Atoms', 'Nitrogen Atoms', 'Oxygen Atoms', 'Carbon Residue Details',
'Nitrogen Residue Details', 'Oxygen Residue Details'])

        # Iterate over PDB files in the directory
        for pdb_file in os.listdir(pdb_dir):
            if pdb_file.endswith('.pdb'):
                pdb_path = os.path.join(pdb_dir, pdb_file)
                counts, details = process_pdb_file(pdb_path)
                if counts:
                    writer.writerow([
                        pdb_file,
                        counts.get('C', 0),
                        counts.get('N', 0),
                        counts.get('O', 0),
                        details.get('Carbon', []),
                        details.get('Nitrogen', []),
                        details.get('Oxygen', [])
                    ])

    print(f"Results have been written to {output_csv}.")
```

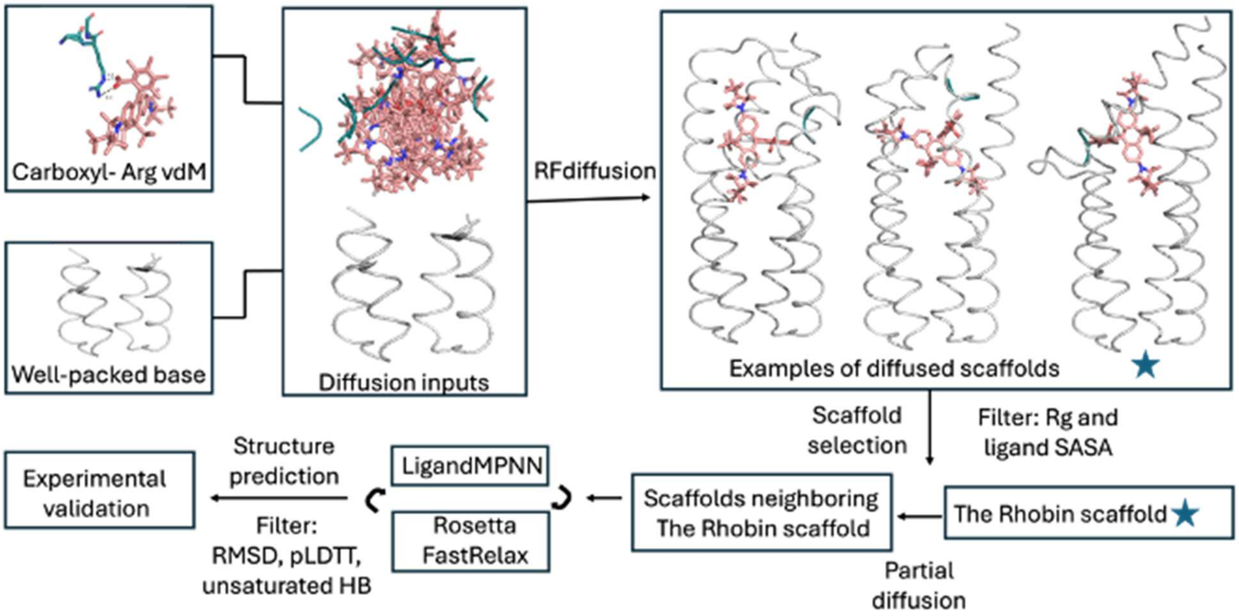


**Fig. S1. Workflow of protein design.**

To design rhodamine-binding proteins, we first used knowledge-guided RFdiffusion to generate rhodamine-specific scaffolds. Three key inputs were provided: (1) a conserved folding core from previously designed ligand-binding helical bundles, (2) an expanded rhodamine template, and (3) a statistically favored arginine positioned to interact with the ligand's carboxylate group. From the diffusion outputs of nine inputs, the Rhobin scaffold (blue star) was selected based on computational filtering and manual evaluation. The selected scaffold then underwent sequence design through iterative cycles of LigandMPNN and Rosetta FastRelax. Designed sequences were filtered by structure prediction, and selected candidates were advanced for experimental validation.

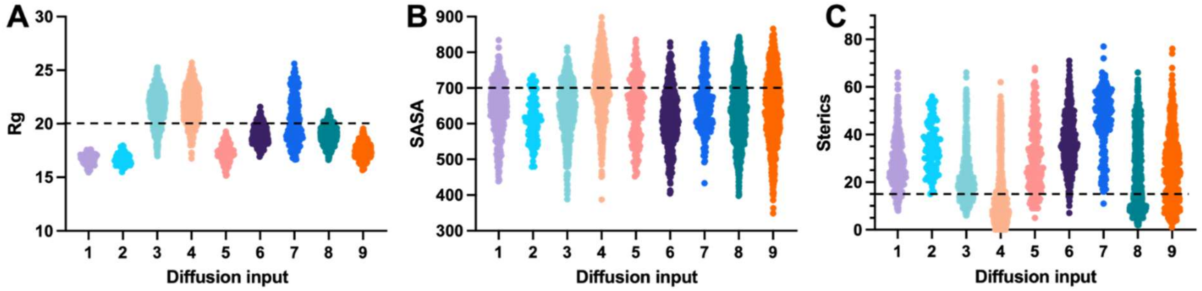


**Fig. S2. Selection of the rhodamine-specific scaffold.**

Statistics of diffused backbones with different starting poses on radius of gyration (Rg, **A**), solvent-accessible surface area (SASA, **B**), and sterics (atoms within 3.5 A of expanded rhodamine template, **C**). The thresholds of these three computational filters were shown as dashed lines, and the ones with lower values for all three metrics were subjected to manual evaluation.

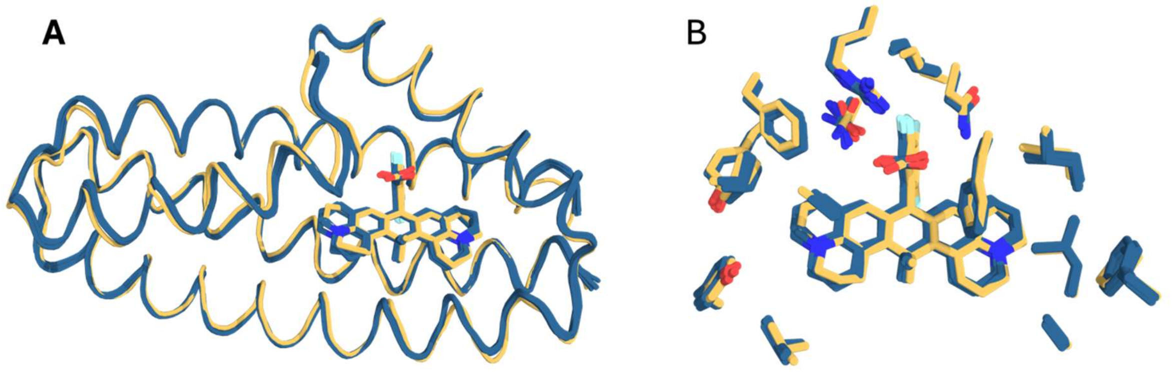


**Fig. S3. The agreement of Rhobin9 design structure with its Chai1 predicted structures.**
(**A**) Five predicted Rhobin9 structures (blue cartoon) complexed with $JF_{660}$ (blue sticks) were aligned to its designed Rhobin9-$JF_{660}$ complex, shown in yellow cartoon and sticks. (**B**) Zoomed-in view of the sidechain structures of the active site in (A).

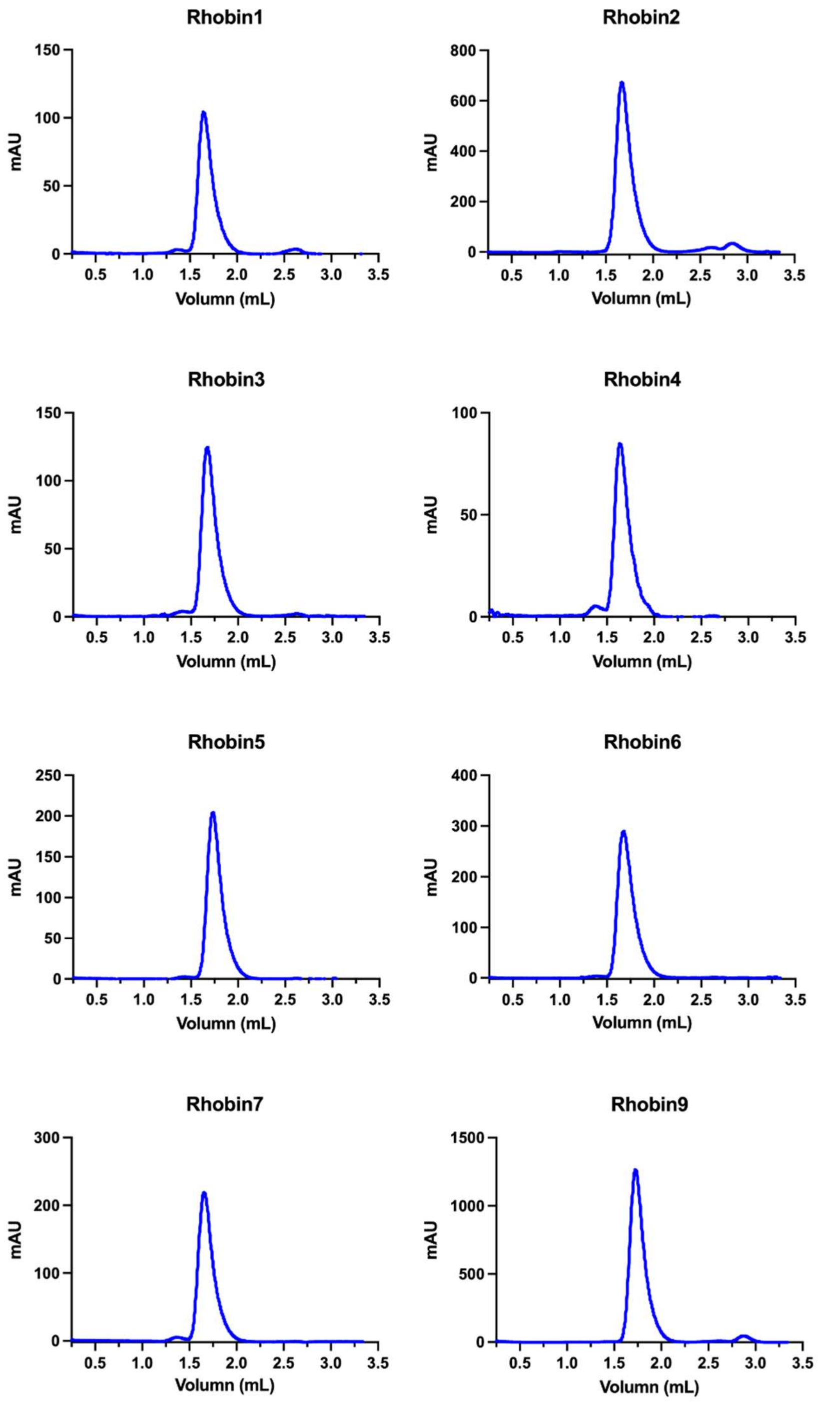


**Fig. S4. FPLC characterization of Rhobin.**

The monomeric state of each Rhobin with binding interaction towards $JF_{660}$ was confirmed with FPLC equipped with Superdex 75 Increase 5/150 analytical column. Protein was monitored with absorbance at 280 nm.

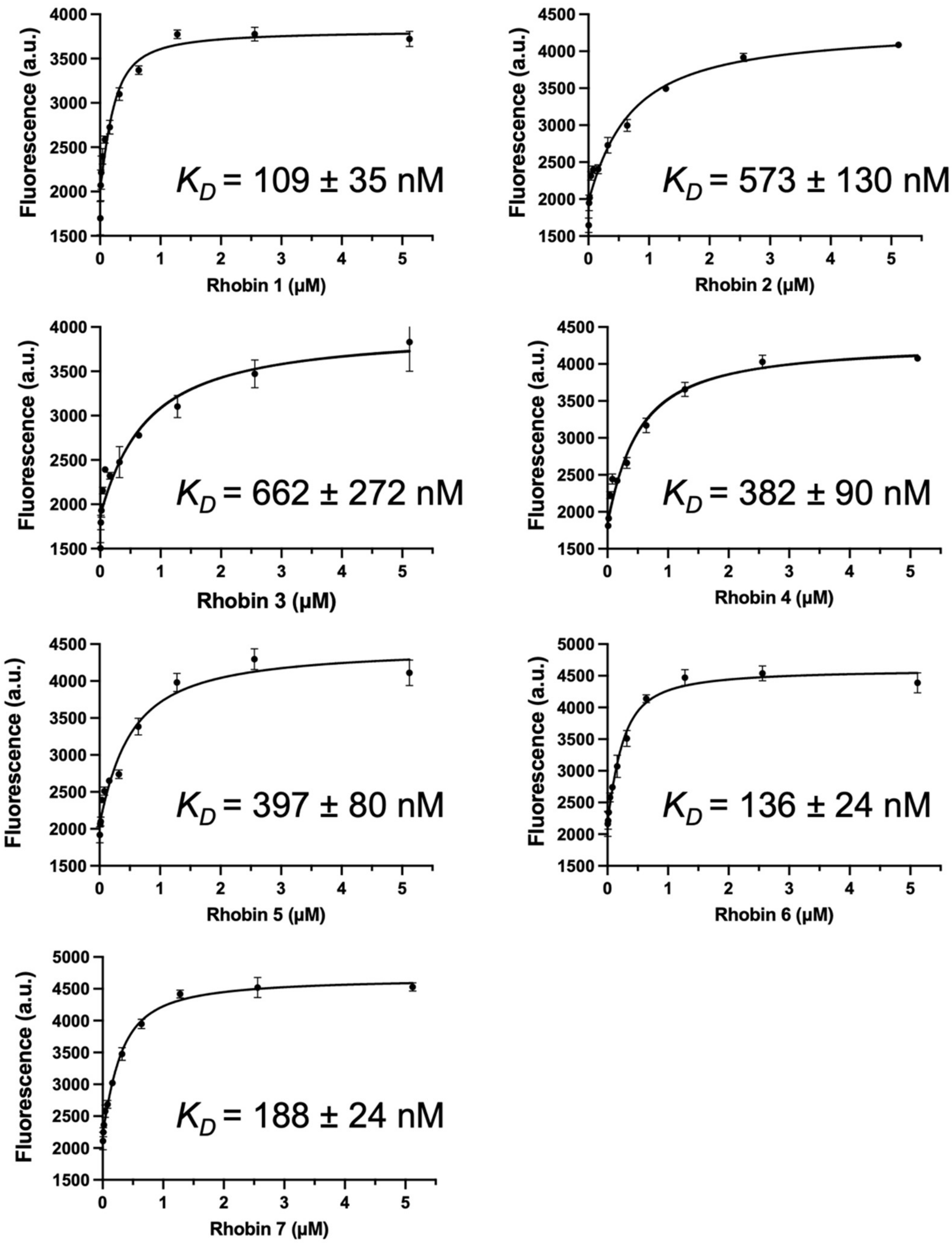


**Fig. S5. Binding constant determination for Rhobin with JF660.**
Binding constants ($K_D$) of Rhobin to $JF_{660}$, measured by fluorescence titration of Rhobin into 200 nM $JF_{660}$. $K_D$ is shown as the mean and standard error of the mean.

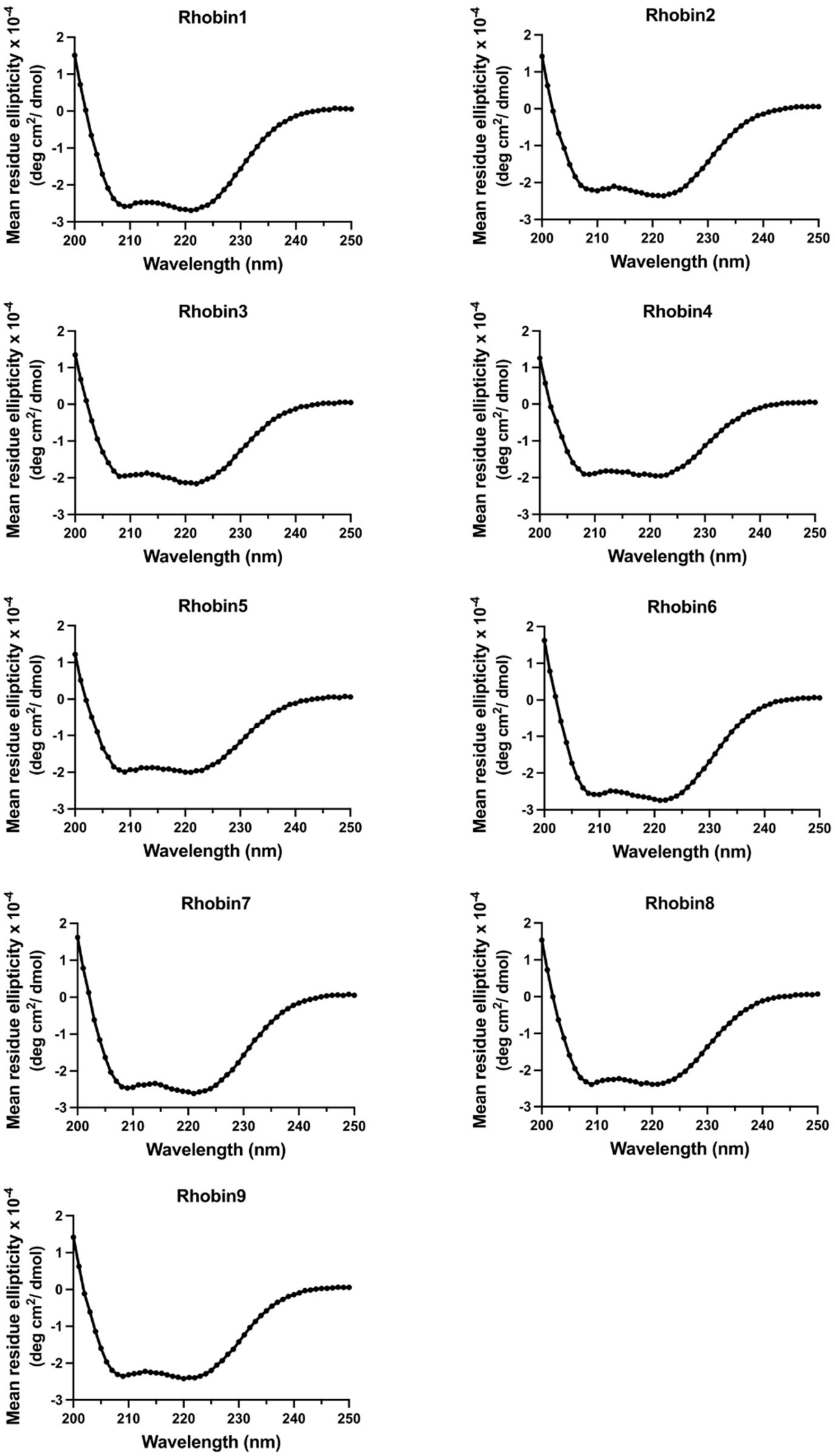


**Fig. S6. Helical structures of Rhobin shown from CD spectrum.**
Proteins were prepared in 10 μM concentrations in PBS buffer. CD spectra were collected on a Jasco J-810 CD spectrometer in a 0.1 cm path-length quartz cuvette.

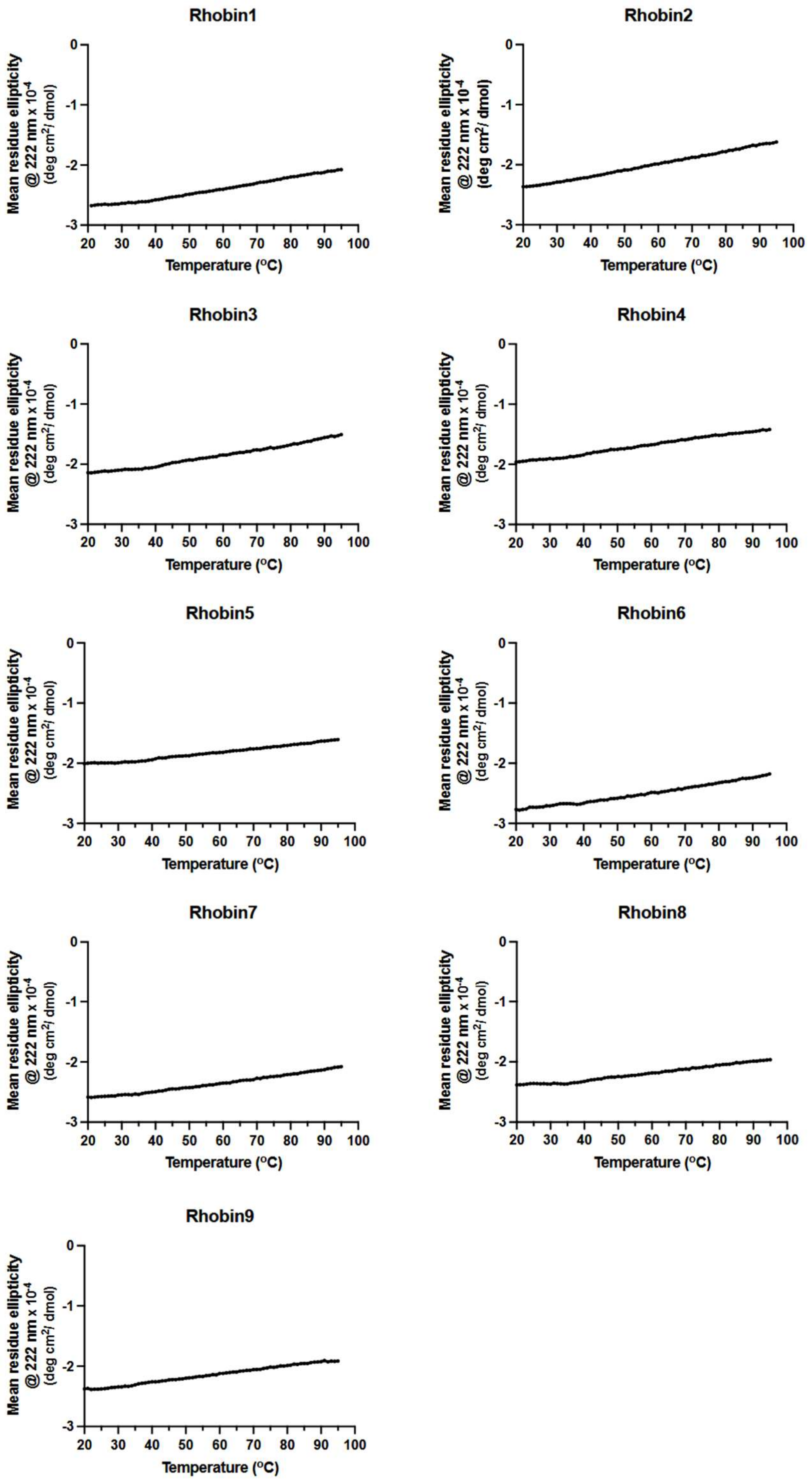


**Fig. S7. Thermostability of Rhobin.**
Proteins were prepared in 10 μM concentrations in PBS buffer. CD spectra were collected on a Jasco J-810 CD spectrometer in a 0.1 cm path-length quartz cuvette. Temperature-dependent data were collected at 222 nm from 20 to 95 °C with an interval of 5 °C and an increase rate of 2 °C/minute.

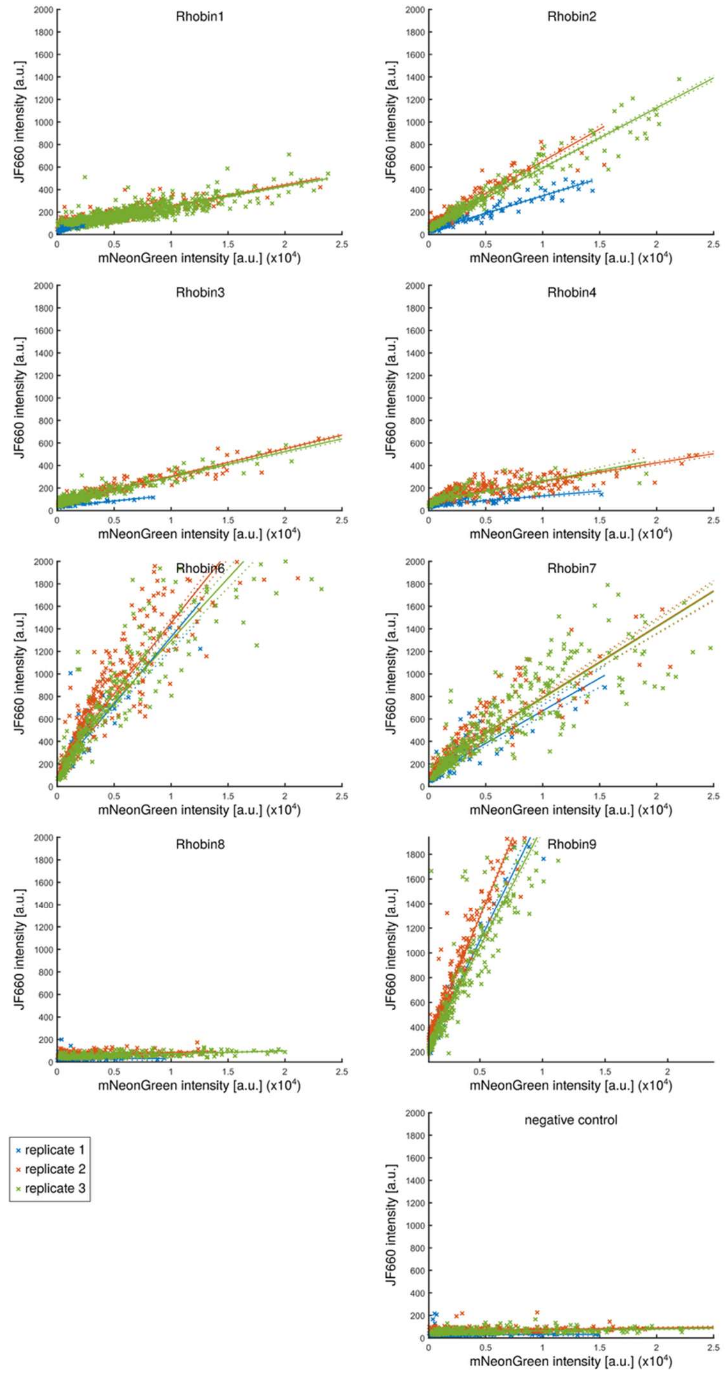


**Fig. S8. Normalized brightness measurements for $JF_{660}$ binding to Rhobin.**

Signal generated through $JF_{660}$ binding to Rhobin variants (Rhobin1-9) and the non-binding frFAST negative control scales linearly with expression level, showing per-cell mean nuclear $JF_{660}$ and mNeonGreen intensities and linear fits for individual replicates.

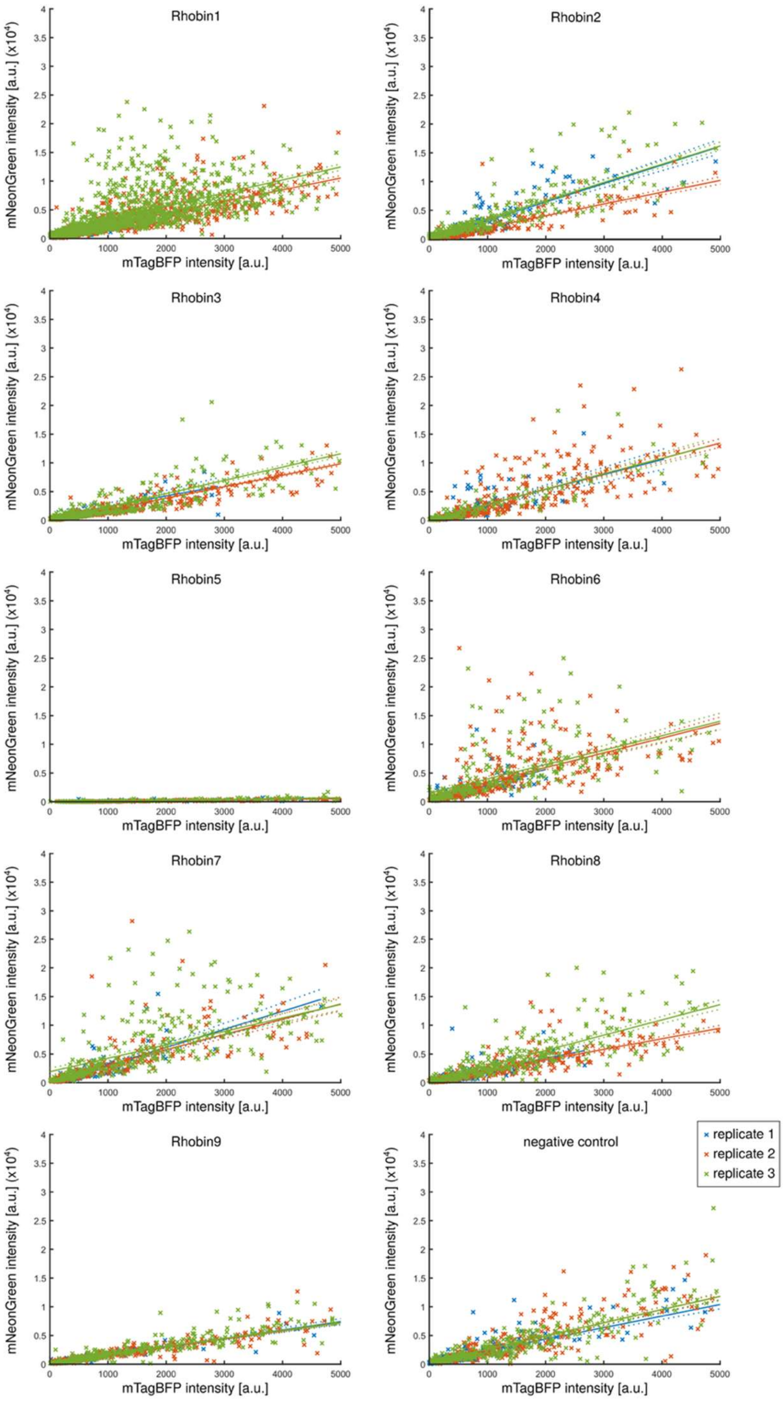


**Fig. S9. Normalized expression of Rhobin variants.**

Fluorescence signal generated from expression reporter mNeonGreen fused at the C-termini of Rhobin variants and negative control. mNeonGreen fluorescence intensity scales linearly with mTagBFP expression level, showing per-cell mean nuclear mNeonGreen and mTagBFP intensities and linear fits from three independent replicates.

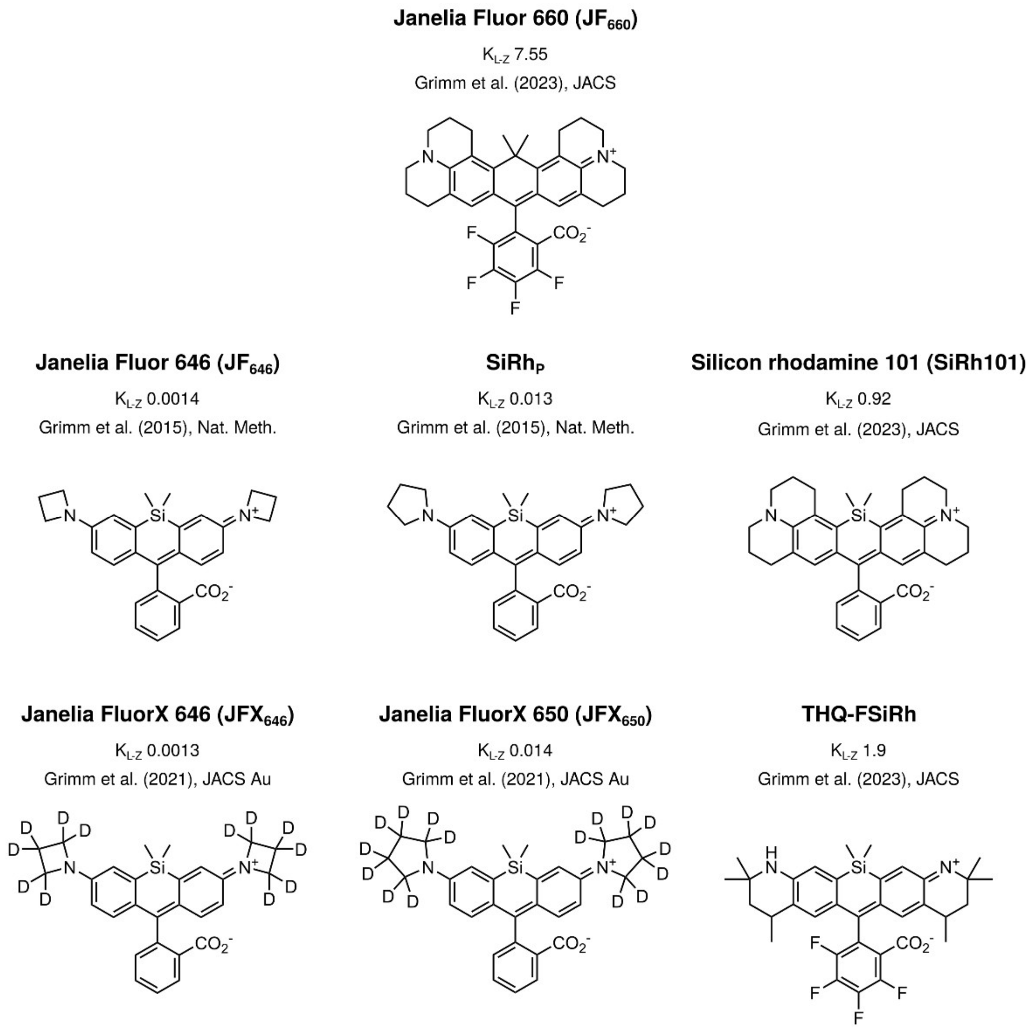


**Fig. S10. Rhodamine dye chemical structures used in this study.**

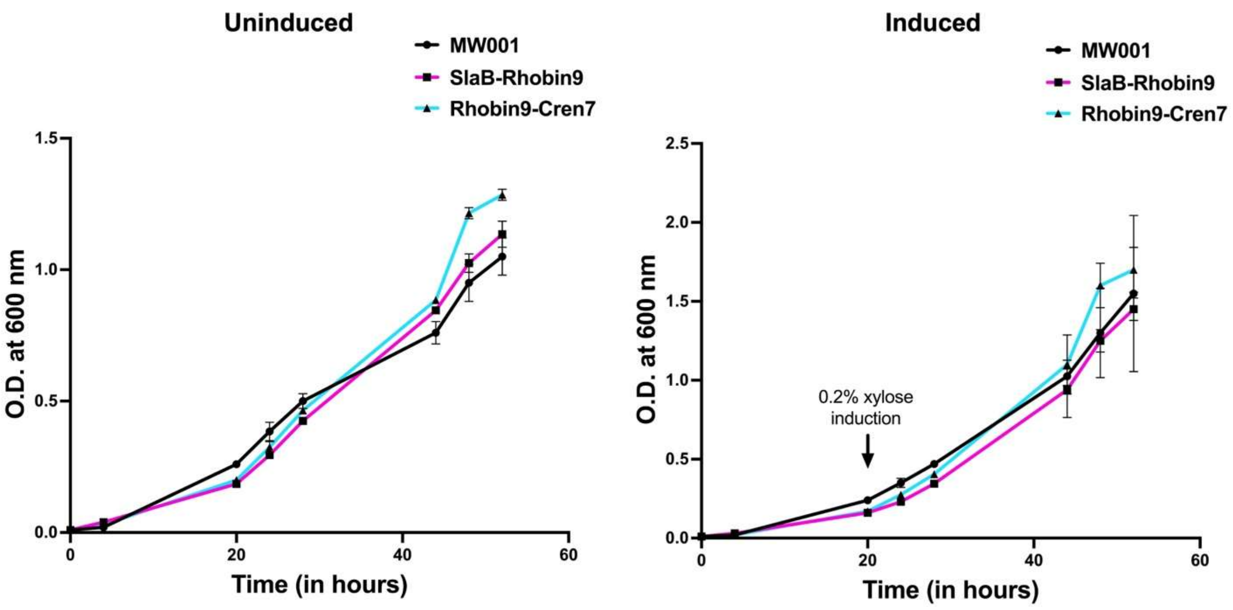


**Fig. S11. Growth comparison of *S. acidocaldarius* cells expressing SlaB-Rhobin9 and Rhobin9-Cren7 with MW001 (WT).**

The left panel shows uninduced cells, while the right panel represents cells induced with 0.2% xylose. Each data point represents the mean, and the error bars indicate the standard deviation (SD) from two biological replicates.

**Table S1. RMSD (Å) of Rhobin designs with their predicted complex structures using Chai1 or apo structures using AF3, including Cα for all residues and all heavy atoms at the active site.**

| | **Chai1** | **AF3** |
|---|---|---|
| Rhobin1 | 0.89 ± 0.06 | 0.87 ± 0.11 |
| Rhobin2 | 0.65 ± 0.05 | 0.75 ± 0.07 |
| Rhobin3 | 0.71 ± 0.05 | 0.80 ± 0.05 |
| Rhobin4 | 0.85 ± 0.04 | 0.82 ± 0.08 |
| Rhobin5 | 0.64 ± 0.05 | 0.81 ± 0.35 |
| Rhobin6 | 0.48 ± 0.03 | 0.56 ± 0.06 |
| Rhobin7 | 0.42 ± 0.03 | 0.51 ± 0.05 |
| Rhobin8 | 0.90 ± 0.02 | 0.82 ± 0.03 |
| Rhobin9 | 0.77 ± 0.03 | 0.85 ± 0.36 |

**Table S2. Numerical values for brightness analysis of binder/fluorophore pairs tested in this study.**

| Fig. | Binder | Fluorophore | Concentration | # exp. | # cells | mean normalized brightness | SD |
|---|---|---|---|---|---|---|---|
| 2D | Rhobin1 | $JF_{660}$ | 100 nM | 3 | 6009 | 0.0187 | 0.0005 |
| 2D | Rhobin2 | $JF_{660}$ | 100 nM | 3 | 2520 | 0.0472 | 0.0142 |
| 2D | Rhobin3 | $JF_{660}$ | 100 nM | 3 | 3186 | 0.0191 | 0.0077 |
| 2D | Rhobin4 | $JF_{660}$ | 100 nM | 3 | 2067 | 0.0145 | 0.0055 |
| 2D | Rhobin6 | $JF_{660}$ | 100 nM | 3 | 2448 | 0.1186 | 0.008 |
| 2D | Rhobin7 | $JF_{660}$ | 100 nM | 3 | 2139 | 0.0615 | 0.003 |
| 2D | Rhobin8 | $JF_{660}$ | 100 nM | 3 | 2427 | 0.0017 | 0.0015 |
| 2D | Rhobin9 | $JF_{660}$ | 100 nM | 3 | 2520 | 0.2155 | 0.0253 |
| 2D | neg. ctrl. | $JF_{660}$ | 100 nM | 3 | 2688 | 0.001 | 0.0007 |
| 2G | Rhobin1 | $JFX_{646}$ | 1 uM | 3 | 1717 | 0.0004 | 0.0002 |
| 2G | Rhobin2 | $JFX_{646}$ | 1 uM | 4 | 1944 | 0.0048 | 0.0015 |
| 2G | Rhobin3 | $JFX_{646}$ | 1 uM | 4 | 1705 | 0.002 | 0.0009 |
| 2G | Rhobin4 | $JFX_{646}$ | 1 uM | 3 | 1081 | 0.0003 | 0.0002 |
| 2G | Rhobin6 | $JFX_{646}$ | 1 uM | 4 | 2441 | 0.0012 | 0.0005 |
| 2G | Rhobin7 | $JFX_{646}$ | 1 uM | 4 | 3067 | 0.0003 | 0.0002 |
| 2G | Rhobin8 | $JFX_{646}$ | 1 uM | 3 | 1576 | 0.0001 | 0 |
| 2G | Rhobin9 | $JFX_{646}$ | 1 uM | 4 | 2623 | 0.0052 | 0.0028 |
| 2G | neg. ctrl. | $JFX_{646}$ | 1 uM | 5 | 3697 | 0.0001 | 0 |
| 2G | Rhobin1 | $JF_{646}$ | 1 uM | 3 | 1781 | 0.0001 | 0.0001 |
| 2G | Rhobin2 | $JF_{646}$ | 1 uM | 3 | 1283 | 0.0019 | 0.0011 |
| 2G | Rhobin3 | $JF_{646}$ | 1 uM | 4 | 3444 | 0.0004 | 0.0002 |
| 2G | Rhobin4 | $JF_{646}$ | 1 uM | 2 | 381 | 0 | 0 |
| 2G | Rhobin6 | $JF_{646}$ | 1 uM | 3 | 1764 | 0.0007 | 0.0005 |
| 2G | Rhobin7 | $JF_{646}$ | 1 uM | 3 | 2236 | 0.0001 | 0.0001 |
| 2G | Rhobin8 | $JF_{646}$ | 1 uM | 3 | 1872 | 0 | 0 |
| 2G | Rhobin9 | $JF_{646}$ | 1 uM | 3 | 1636 | 0.0019 | 0.002 |
| 2G | neg. ctrl. | $JF_{646}$ | 1 uM | 5 | 3790 | 0 | 0 |
| 2G | Rhobin1 | $JFX_{650}$ | 1 uM | 3 | 3172 | 0.0028 | 0.0001 |
| 2G | Rhobin2 | $JFX_{650}$ | 1 uM | 4 | 3944 | 0.0652 | 0.0242 |
| 2G | Rhobin3 | $JFX_{650}$ | 1 uM | 3 | 1126 | 0.0512 | 0.0054 |
| 2G | Rhobin4 | $JFX_{650}$ | 1 uM | 2 | 731 | 0.0006 | 0.0004 |
| 2G | Rhobin6 | $JFX_{650}$ | 1 uM | 2 | 1247 | 0.0097 | 0.0053 |
| 2G | Rhobin7 | $JFX_{650}$ | 1 uM | 3 | 2606 | 0.0041 | 0.0017 |
| 2G | Rhobin8 | $JFX_{650}$ | 1 uM | 3 | 2240 | 0.0006 | 0.0002 |
| 2G | Rhobin9 | $JFX_{650}$ | 1 uM | 4 | 3344 | 0.0538 | 0.0212 |
| 2G | neg. ctrl. | $JFX_{650}$ | 1 uM | 3 | 1944 | 0.0004 | 0.0001 |
| 2G | Rhobin1 | $SiRh_P$ | 1 uM | 3 | 1812 | 0.0011 | 0.0003 |
| 2G | Rhobin2 | $SiRh_P$ | 1 uM | 4 | 2673 | 0.0479 | 0.0152 |
| 2G | Rhobin3 | $SiRh_P$ | 1 uM | 5 | 1532 | 0.0286 | 0.0106 |

| | | | | | | | |
|---|---|---|---|---|---|---|---|
| 2G | Rhobin4 | SiRh$_{P}$ | 1 uM | 2 | 481 | 0.0007 | 0.0001 |
| 2G | Rhobin6 | SiRh$_{P}$ | 1 uM | 3 | 982 | 0.0104 | 0.0035 |
| 2G | Rhobin7 | SiRh$_{P}$ | 1 uM | 3 | 1389 | 0.0036 | 0.0013 |
| 2G | Rhobin8 | SiRh$_{P}$ | 1 uM | 2 | 488 | 0.0003 | 0.0001 |
| 2G | Rhobin9 | SiRh$_{P}$ | 1 uM | 4 | 1752 | 0.0368 | 0.0128 |
| 2G | neg. ctrl. | SiRh$_{P}$ | 1 uM | 5 | 2711 | 0.0002 | 0.0001 |
| 2G | Rhobin1 | THQ-FSiRh | 1 uM | 3 | 2069 | 0.013 | 0.0093 |
| 2G | Rhobin2 | THQ-FSiRh | 1 uM | 3 | 793 | 0.0175 | 0.0191 |
| 2G | Rhobin3 | THQ-FSiRh | 1 uM | 3 | 670 | 0.0074 | 0.0047 |
| 2G | Rhobin4 | THQ-FSiRh | 1 uM | 3 | 446 | 0.02 | 0.015 |
| 2G | Rhobin6 | THQ-FSiRh | 1 uM | 4 | 1083 | 0.049 | 0.0151 |
| 2G | Rhobin7 | THQ-FSiRh | 1 uM | 3 | 789 | 0.0214 | 0.014 |
| 2G | Rhobin8 | THQ-FSiRh | 1 uM | 3 | 900 | -0.0003 | 0.0141 |
| 2G | Rhobin9 | THQ-FSiRh | 1 uM | 4 | 1311 | 0.0331 | 0.0201 |
| 2G | neg. ctrl. | THQ-FSiRh | 1 uM | 6 | 3153 | 0.0077 | 0.0065 |
| 2G | Rhobin1 | SiRh101 | 1 uM | 3 | 1614 | 0.0016 | 0.001 |
| 2G | Rhobin2 | SiRh101 | 1 uM | 3 | 825 | 0.0086 | 0.0042 |
| 2G | Rhobin3 | SiRh101 | 1 uM | 4 | 1212 | 0.0051 | 0.003 |
| 2G | Rhobin4 | SiRh101 | 1 uM | 3 | 551 | 0.0021 | 0.0005 |
| 2G | Rhobin6 | SiRh101 | 1 uM | 4 | 1607 | 0.0249 | 0.0108 |
| 2G | Rhobin7 | SiRh101 | 1 uM | 4 | 2106 | 0.0077 | 0.0034 |
| 2G | Rhobin8 | SiRh101 | 1 uM | 4 | 1354 | 0.001 | 0.0005 |
| 2G | Rhobin9 | SiRh101 | 1 uM | 5 | 2406 | 0.0335 | 0.0214 |
| 2G | neg. ctrl. | SiRh101 | 1 uM | 6 | 3546 | 0.0006 | 0.0004 |
| 2H | HaloTag7 | Halo-JF$_{660}$ | 100 nM | 4 | 775 | 0.0026 | 0.0018 |
| 2H | reHaloF | Halo-JF$_{660}$ | 100 nM | 3 | 1127 | 0.0001 | 0.0001 |
| 2H | Rhobin2 | JF$_{660}$ | 100 nM | 4 | 3779 | 0.0572 | 0.023 |
| 2H | Rhobin9 | JF$_{660}$ | 100 nM | 4 | 745 | 0.2171 | 0.0203 |
| 2H | neg. ctrl | Halo-JF$_{660}$ | 100 nM | 4 | 1706 | 0.0001 | 0 |
| 2H | neg. ctrl | JF$_{660}$ | 100 nM | 3 | 1020 | 0.001 | 0.0008 |
| 2H | HaloTag7 | Halo-JFX$_{650}$ | 100 nM | 4 | 1981 | 0.2683 | 0.0186 |
| 2H | reHaloF | Halo-JFX$_{650}$ | 100 nM | 5 | 964 | 0.0242 | 0.0094 |
| 2H | Rhobin2 | JFX$_{650}$ | 100 nM | 4 | 1330 | 0.032 | 0.0026 |
| 2H | Rhobin9 | JFX$_{650}$ | 100 nM | 4 | 2947 | 0.0193 | 0.0031 |
| 2H | neg. ctrl | Halo-JFX$_{650}$ | 100 nM | 4 | 2925 | 0 | 0 |
| 2H | neg. ctrl | JFX$_{650}$ | 100 nM | 3 | 2709 | 0.0001 | 0 |
| 2H | frFAST | HPAR-3OM | 5 uM | 4 | 3113 | 0.0008 | 0 |
| 2H | neg. ctrl | HPAR-3OM | 5 uM | 3 | 3139 | 0 | 0 |
| 3C | Rhobin2 | SNAP-JFX$_{650}$ | 1 uM | 3 | 2335 | 0.0037 | 0.0002 |
| 3C | Rhobin2 | Halo-JFX$_{650}$ | 1 uM | 3 | 2654 | 0.0007 | 0 |
| 3C | Rhobin2 | JFX$_{650}$ | 1 uM | 3 | 3127 | 0.0347 | 0.0017 |
| 3C | neg. ctrl | JFX$_{650}$ | 1 uM | 3 | 3090 | 0.0004 | 0 |

**Table S3. Experimental parameters for microscopy datasets.**

| Fig | Sample | Construct(s) | Dye | Dye conc | Microscope | Condition |
|---|---|---|---|---|---|---|
| 2B | U2OS | 19, 39 | JF$_{660}$ | 100 nM | Widefield | no-wash |
| 2C | U2OS | 19, 39 | JF$_{660}$ | 100 nM | Widefield | no-wash |
| 2D | U2OS | 10-14, 16-19 | | | Widefield | no-wash |
| 2E | U2OS | 10-19 | - | - | Widefield | no-wash |
| 2G | U2OS | 10-14, 16-19, 39 | JFX$_{646}$<br>JF$_{646}$<br>JFX$_{650}$<br>SiRh$_{P}$<br>THQ-FSiRh<br>SiRh101 | 1µM | Widefield | no-wash |
| 2H | U2OS | 12, 19, 37, 38, 39 | Halo-JF$_{660}$<br>Halo-JFX$_{650}$<br>JF$_{660}$<br>JFX$_{650}$<br>HPAR-30M | 100 nM<br>100 nM<br>100 nM<br>100 nM<br>5 µM | Widefield | no-wash |
| 3B | U2OS LP | lines 06-15 | SiRh$_{P}$ | 50 nM | Spinning-disk | no-wash |
| 3C | U2OS | 12 | Halo-JFX$_{650}$<br>SNAP-JFX$_{650}$<br>JFX$_{650}$ | 1 µM | Widefield | no-wash |
| 3D | U2OS | 31, 36, 37 | Halo-JF$_{503}$<br>SNAP-JF$_{571}$<br>SiRh$_{P}$ | 100 nM<br>1 µM<br>100 nM | Spinning-disk | post-wash/<br>no-wash* |
| 4B,C,D | U2OS | 35 | JFX$_{650}$ | 100 nM | STED | no-wash |
| 4E | U2OS | line 13 | SiRh$_{P}$ | 2 µM | STED | no-wash |
| 4F | U2OS LP | line 13 | SiRh$_{P}$ | 2 µM | STED | no-wash |
| 4G | U2OS | 37 | Halo-JFX$_{650}$ | 1 µM | STED | no-wash |
| 4I,J,K | U2OS | 33, 37, 38 | SiRh$_{P}$ | 5 nM | TIRF | no-wash |
| 5B | *S. acidocaldarius* | wt, 43, 44 | JF$_{660}$ | 250 nM | iSIM | Post-wash |
| 5C | *S. acidocaldarius* | 43, 44 | JF$_{660}$ | 250 nM | iSIM | Post-wash |

* Samples were washed post-staining with HaloTag and SNAP-tag ligands. No-wash imaging with Rhobin2 ligand.

### Table S4. List of plasmids used in this study.

| ID | Name | Purpose | Addgene ID (if available) | Source/Reference |
|---|---|---|---|---|
| 01 | pET29b-Rhobin1 | Bacterial expression of Rhobin1 | | This study |
| 02 | pET29b-Rhobin2 | Bacterial expression of Rhobin2 | 239018 | This study |
| 03 | pET29b-Rhobin3 | Bacterial expression of Rhobin3 | | This study |
| 04 | pET29b-Rhobin4 | Bacterial expression of Rhobin4 | | This study |
| 05 | pET29b-Rhobin5 | Bacterial expression of Rhobin5 | | This study |
| 06 | pET29b-Rhobin6 | Bacterial expression of Rhobin6 | | This study |
| 07 | pET29b-Rhobin7 | Bacterial expression of Rhobin7 | | This study |
| 08 | pET29b-Rhobin8 | Bacterial expression of Rhobin8 | | This study |
| 09 | pET29b-Rhobin9 | Bacterial expression of Rhobin9 | 239019 | This study |
| 10 | pHaloTag-H2B-mNeonGreen3K-IREs-NLS-mTagBFP-NLSx2 | Mammalian expression of HaloTag7-H2B with an expression marker NLS-mTagBFP-NLSx2 | | This study |
| 11 | pRhobin1-H2B-mNeonGreen3K-IRES-NLS-mTagBFP-NLSx2 | Mammalian expression of Rhobin1-H2B with an expression marker NLS-mTagBFP-NLSx2 | | This study |
| 12 | pRhobin2-H2B-mNeonGreen3K-IRES-NLS-mTagBFP-NLSx2 | Mammalian expression of Rhobin2-H2B with an expression marker NLS-mTagBFP-NLSx2 | 239020 | This study |
| 13 | pRhobin3-H2B-mNeonGreen3K-IRES-NLS-mTagBFP-NLSx2 | Mammalian expression of Rhobin3-H2B with an expression marker NLS-mTagBFP-NLSx2 | | This study |
| 14 | pRhobin4-H2B-mNeonGreen3K-IRES-NLS-mTagBFP-NLSx2 | Mammalian expression of Rhobin4-H2B with an expression marker NLS-mTagBFP-NLSx2 | | This study |
| 15 | pRhobin5-H2B-mNeonGreen3K-IRES-NLS-mTagBFP-NLSx2 | Mammalian expression of Rhobin5-H2B with an expression marker NLS-mTagBFP-NLSx2 | | This study |
| 16 | pRhobin6-H2B-mNeonGreen3K-IRES-NLS-mTagBFP-NLSx2 | Mammalian expression of Rhobin6-H2B with an expression marker NLS-mTagBFP-NLSx2 | | This study |
| 17 | pRhobin7-H2B-mNeonGreen3K-IRES-NLS-mTagBFP-NLSx2 | Mammalian expression of Rhobin7-H2B with an expression marker NLS-mTagBFP-NLSx2 | | This study |
| 18 | pRhobin8-H2B-mNeonGreen3K-IRES-NLS-mTagBFP-NLSx2 | Mammalian expression of Rhobin8-H2B with an expression marker NLS-mTagBFP-NLSx2 | | This study |
| 19 | pRhobin9-H2B-mNeonGreen3K-IRES-NLS-mTagBFP-NLSx2 | Mammalian expression of Rhobin9-H2B with an expression marker NLS-mTagBFP-NLSx2 | 239021 | This study |
| 20 | pPa01attP-pCMV-H2B-Rhobin2-IRES-NLS-mTagBFP-NLSx2 | Donor plasmid for U2OS H2B-Rhobin2 stable cell line generation | | This study |
| 21 | pPa01attP-pCMV-NPM1-Rhobin2-IRES-NLS-mTagBFP-NLSx2 | Donor plasmid for U2OS NPM1-Rhobin2 stable cell line generation | | This study |
| 22 | pPa01attP-pCMV-FBL-Rhobin2-IRES-NLS-mTagBFP-NLSx2 | Donor plasmid for U2OS FBL-Rhobin2 stable cell line generation | | This study |
| 23 | pPa01attP-pCMV-Rhobin2-LMNA-IRES-NLS-mTagBFP-NLSx2 | Donor plasmid for U2OS Rhobin2-LMNA stable cell line generation | | This study |
| 24 | pPa01attP-pCMV-LifeAct-Rhobin2-IRES-NLS-mTagBFP-NLSx2 | Donor plasmid for U2OS LifeAct-Rhobin2 stable cell line generation | | This study |
| 25 | pPa01attP-pCMV-Rhobin2-Rab5a-IRES-NLS-mTagBFP-NLSx2 | Donor plasmid for U2OS Rhobin2-Rab5a stable cell line generation | | This study |
| 26 | pPa01attP-pCMV-Rhobin2-CLTB-IRES-NLS-mTagBFP-NLSx2 | Donor plasmid for U2OS Rhobin2-CLTB stable cell line generation | | This study |
| 27 | pPa01attP-pCMV-Rhobin2-SEC61B-IRES-NLS-mTagBFP-NLSx2 | Donor plasmid for U2OS Rhobin2-SEC61B stable cell line generation | | This study |
| 28 | pPa01attP-pCMV-Rhobin2-Giantin-IRES-NLS-mTagBFP-NLSx2 | Donor plasmid for U2OS Rhobin2-Giantin stable cell line generation | | This study |
| 29 | pPa01attP-pCMV-TOMM20-Rhobin2-IRES-NLS-mTagBFP-NLSx2 | Donor plasmid for U2OS TOMM20-Rhobin2 stable cell line generation | | This study |
| 30 | pEF1a-Pa01 | Mammalian expression Pa01 integrase | | This study, EF1a-Pa01 cDNA from |

| | | | | |
|---|---|---|---|---|
| | | | | Addgene # 194360)(39) |
| 31 | pTOMM20-Rhobin2-IRES-NLS-mTagBFP-NLSx2 | Mammalian expression of TOMM20-Rhobin2 with an expression marker NLS-mTagBFP-NLSx2 | 239022 | This study |
| 32 | pTOMM20-Rhobin9-IRES-NLS-mTagBFP-NLSx2 | Mammalian expression of TOMM20-Rhobin9 with an expression marker NLS-mTagBFP-NLSx2 | 239023 | This study |
| 33 | pRhobin2-SEC61B-IRES-NLS-mTagBFP-NLSx2 | Mammalian expression of Rhobin2-SEC61B with an expression marker NLS-mTagBFP-NLSx2 | 239024 | This study |
| 34 | pRhobin9-SEC61B-IRES-NLS-mTagBFP-NLSx2 | Mammalian expression of Rhobin9-SEC61B with an expression marker NLS-mTagBFP-NLSx2 | 239025 | This study |
| 35 | pLifeAct-Rhobin2-IRES-NLS-mTagBFP-NLSx2 | Mammalian expression of LifeAct-Rhobin2 with an expression marker NLS-mTagBFP-NLSx2 | | This study |
| 36 | pSNAPf-H2B | Mammalian expression of H2B-SNAPf | 101124 | Unpublished (Gift from New England Biolabs & Ana Egana) |
| 37 | pHaloTag7-SEC61B-IRES-NLS-mTagBFP-NLSx2 | Mammalian expression of HaloTag7-SEC61B with an expression marker NLS-mTagBFP-NLSx2 | | This study |
| 38 | reHaloF-SEC61B-IRES-NLS-mTagBFP-NLSx2 | Mammalian expression of reHaloF-SEC61B with an expression marker NLS-mTagBFP-NLSx2 | | This study |
| 39 | frFAST-H2B-mNeoNGreen-IRES-NLS-mTagBFP-NLSx2 | Mammalian expression of frFAST-H2B-mNeonGreen with an expression marker NLS-mTagBFP-NLSx2 | | This study |
| 40 | Sniper2L | Transient expression of Sniper2L Cas9 for Pa01 attB landing pad site integration into *AAVS1* locus | 193856 | (*55*) |
| 41 | pU6-AAVS1-HA-sgRNA | Expression of single guide RNA targeting *AAVS1* locus | | (*48*) |
| 42 | AAVS1-Pa01attB-BFP-Puro-HA | Donor plasmid for integration of Pa01 attB followed TagBFP-NLS-T2A-Puro | | (*48*) |
| 43 | pACO48 | Archaeal expression of SlaB- GGTGGGGSGG -Rhobin9 | | This study |
| 44 | pACO49 | Archaeal expression of Rhobin9-GGTGGGGSGG-Cren7 | | This study |

**Table S5. List of plasmids used in this study.**

| ID | Name | Description | Source/Reference |
|---|---|---|---|
| 01 | *E, coli* BL21(DE3) | Bacterial expression | New England Biolabs |
| 02 | *E. coli* HST08 Stellar | Cloning and plasmid amplification | TaKaRa |
| 03 | ER1821 | Methylated plasmid production | Sonja-Verena Albers (University of Freiburg, Germany). |
| 04 | U2OS (WT) | Mammalian expression | ATCC |
| 05 | U2OS LP | Landing Pad site attB-integrated at AAVS1 for stable cell generation. | Huang Lab |
| 06 | U2OS LP H2B-Rhobin2 | Stable expression of H2B-Rhobin2 | This study |
| 07 | U2OS LP NPM1-Rhobin2 | Stable expression of NPM1-Rhobin2 | This study |
| 08 | U2OS LP FBL-Rhobin2 | Stable expression of FBL-Rhobin2 | This study |
| 09 | U2OS LP Rhobin2-LMNA | Stable expression of Rhobin2-LMNA | This study |
| 10 | U2OS LP LifeAct-Rhobin2 | Stable expression of LifeAct-Rhobin2 | This study |
| 11 | U2OS LP Rhobin2-Rab5a | Stable expression of Rhobin2-Rab5a | This study |
| 12 | U2OS LP Rhobin2-CLTB | Stable expression of Rhobin2-CLTB | This study |
| 13 | U2OS LP Rhobin2-SEC61B | Stable expression of Rhobin2-SEC61B | This study |
| 14 | U2OS LP Rhobin2-Giantin | Stable expression of Rhobin2-Giantin | This study |
| 15 | U2OS LP TOMM20-Rhobin2 | Stable expression of TOMM20-Rhobin2 | This study |
| *16* | *S. acidocaldarius* MW001 (WT) | Extremophile *S. acidocaldarius* strain cells for archaeal expression | Sonja-Verena Albers (University of Freiburg, Germany). |

**Table S6. Amino acid sequences of designed proteins.**

| Protein | Amino acid sequence |
|---|---|
| Rhobin1 | SLNQKFIEIYEKLKEYFKKLVETGTKLAAALRAGDRAAAGKLLEEFLKTYE EMVKYGDKEKEKALTILTEEELQAILDQNQEELKKLGVELTNEEVEEMVA ELKRAFEAGDTAAAAAIVEKLVKYYQALIVIAEKQIAKLKSQI |
| Rhobin2 | SEKNQFIIDIYEKLNEYHKKLLELARALAEALRAGDRARAGRLLEEFLATF KAMKAYGDAKEVESFLYFSEEELAAIFAESDVERKKLGVTLKIPEVVAMV EELGRAFEAGDTATAAAIVEKLVKFYEANIIIGQKRIDELKSKI |
| Rhobin3 | SEKNQFIIDIYEKLNEYHKKLLELARALAEALRAGDRARAGRLLEEFLATF KALKAYGDAKEVESFLYFSEEELAAIFAESDVERKKLGVTLKIPEVVAMVE ELGRAFEAGDTATAAAIVEKLVKFYEANIIIGQKRIDELKSKI |
| Rhobin4 | SLLEEDIKILEKVLEYLKKLTELGRRLIEALRAGDRERAGRLLREFLAAYEE MVKYGNEELKKALEIYTEEELQAIFEKNQEELKKLGVTLTNEEVDALVREL GRAFEAGDLAAAAEIAEKLVAFYEALIVIGEKQIKEYKSKI |
| Rhobin5 | SEKKKKLIEIYEKIIGYLKKLYDLARKLAEALRAGDRERAGKLLDEFLAVY NEMVAIGKKMLEEAKKVFTEEEIQKIFEENDVKRKELGVDLNLAEVDRLV AELRAAFEAGDTARAAEIVERLVKFYQANIIIGEERIKKLKSEL |
| Rhobin6 | SEKNKFLIDIYETAIKYLEKMLELAERLAAAMRAGDRATADVLLKEFVAV YEELVKYGEKKKVEAKKVFTEEELKAIFDESDKKRKELGVTLKVPEVDTM VKELVAAFEAGDLETAAAIVEKLVKFYKANIIIGKERIKELKSKL |
| Rhobin7 | SEKNKFLIDIYETLIKYLEKMLELAERLAAAMRAGDRATADVLLKEFVAV YEELVKYGEKKKVEAKKVFTEEELKAIFDESDKKRKELGVTLKVPEVDTM VKELVAAFEAGDLETAAAIVEKLVKFYKANIIIGKERIKELKSKL |
| Rhobin8 | SKNQYIIDILEKLIKYHEKLLKLGRELAEALRAGDRERAGKLLEEYLAVYE EMVKYGEKQKETAKEVFSEEELQKIFEESVKELKKLGVEMTNEEVNQAVK ELRAAFEAGDTARAAAIVEKLVKYYQALIIIGNKRIKELKSKL |
| Rhobin9 | SEKKKKLIEIYEKIIGYLKKLYDLARKLAEALRAGDRERAGKLLDEFLAVY NEMVAIGKKMLEEAKKVFTEEEIQKIFEENDVKRKELGVDLNAAEVDRLV AELRAAFEAGDTARAAEIVERLVKFYQANIIIGEERIKKLKSEL |

**Movie S1. Live-cell STED microscopy of LifeAct-Rhobin2 in U2OS cells.**

U2OS cells transiently expressing LifeAct-Rhobin2 were labeled with 100 nM $JFX_{650}$ and imaged with 2D STED depletion pattern and TauSTED gating at 82 sec per frame.

**Movie S2. Live-cell STED microscopy of Rhobin2-SEC61B in U2OS cells.**

U2OS cells transiently expressing Rhobin2-SEC61B were labeled with 2 μM $SiRh_P$ and imaged with 2D STED depletion pattern and TauSTED gating at 2.58 sec per frame.

**Movie S3. Single-molecule blinking induced by SiRhP binding to Rhobin2.**

U2OS cells transiently expressing Rhobin2-SEC61B were chemically fixed, labeled with 5 nM $SiRh_P$ and imaged under near-TIRF illumination.

**Movie S4. Live-cell imaging of Rhobin9-Cren7 in *S. acidocaldarius*.**

*S. acidocaldarius* cells expressing Rhobin9-Cren7 undergoing cell division and imaged with iSIM at 75 °C.

**Movie S5. Live-cell imaging of SlaB-Rhobin9 in *S. acidocaldarius*.**

*S. acidocaldarius* cells expressing SlaB-Rhobin9 undergoing cell division and imaged with iSIM at 75 °C. Fluorescence images were acquired together with every $6^{th}$ brightfield image.
